# Regulation of Diseases-Associated Microglia in the Optic Nerve by Lipoxin B_4_ and Ocular Hypertension

**DOI:** 10.1101/2024.03.18.585452

**Authors:** Shubham Maurya, Maggie Lin, Shruthi Karnam, Tanirika Singh, Matangi Kumar, Emily Ward, Jeremy Sivak, John G Flanagan, Karsten Gronert

## Abstract

**Background:** The resident astrocyte-retinal ganglion cell (RGC) lipoxin circuit is impaired during retinal stress, which includes ocular hypertension-induced neuropathy. Lipoxin B_4_ produced by homeostatic astrocytes directly acts on RGCs to increase survival and function in ocular hypertension-induced neuropathy. RGC death in the retina and axonal degeneration in the optic nerve are driven by the complex interactions between microglia and macroglia. Whether LXB_4_ neuroprotective actions include regulation of other cell types in the retina and/or optic nerve is an important knowledge gap.

**Methods:** Cellular targets and signaling of LXB_4_ in the retina were defined by single-cell RNA sequencing. Retinal neurodegeneration was induced by injecting silicone oil into the anterior chamber of the mouse eyes, which induced sustained and stable ocular hypertension. Morphological characterization of microglia populations in the retina and optic nerve was established by MorphOMICs and pseudotime trajectory analyses. The pathways and mechanisms of action of LXB_4_ in the optic nerve were investigated using bulk RNA sequencing. Transcriptomics data was validated by qPCR and immunohistochemistry. Differences between experimental groups were assessed by Student’s t-test and one-way ANOVA.

**Results:** Single-cell transcriptomics identified microglia as a primary target for LXB_4_ in the healthy retina. LXB_4_ downregulated genes that drive microglia environmental sensing and reactivity responses. Analysis of microglial function revealed that ocular hypertension induced distinct, temporally defined, and dynamic phenotypes in the retina and, unexpectedly, in the distal myelinated optic nerve. Microglial expression of CD74, a marker of disease-associated microglia in the brain, was only induced in a unique population of optic nerve microglia, but not in the retina. Genetic deletion of lipoxin formation correlated with the presence of a CD74 optic nerve microglia population in normotensive eyes, while LXB_4_ treatment during ocular hypertension shifted optic nerve microglia toward a homeostatic morphology and non-reactive state and downregulated the expression of CD74. Furthermore, we identified a correlation between CD74 and phospho-phosphoinositide 3-kinases (p-PI3K) expression levels in the optic nerve, which was reduced by LXB_4_ treatment.

**Conclusion:** We identified early and dynamic changes in the microglia functional phenotype, reactivity, and induction of a unique CD74 microglia population in the distal optic nerve as key features of ocular hypertension-induced neurodegeneration. Our findings establish microglia regulation as a novel LXB_4_ target in the retina and optic nerve. LXB_4_ maintenance of a homeostatic optic nerve microglia phenotype and inhibition of a disease-associated phenotype are potential neuroprotective mechanisms for the resident LXB_4_ pathway.

## Introduction

Ocular hypertension (OHT) is the primary risk factor responsible for initiating the cascade of events leading to the irreversible retinal neurodegeneration that we know as glaucoma. The only existing therapeutic approach for glaucoma involves the use of topical hypotensive drugs to lower intraocular pressure (IOP)[1–4]. However, glaucoma has a multifactorial etiology, and the precise sequence of events and effector cells that trigger and drive the cascades resulting in retinal ganglion cell (RGC) degeneration[5] are largely unknown. Consequently, there is a pressing need to elucidate these mechanisms, identify new therapeutic targets, and develop treatments to prevent and counteract glaucomatous neurodegeneration.

We recently identified a resident neuroprotective lipid mediator pathway in the retina[6]. Metabolomic screening of homeostatic astrocytes identified lipoxins (LXA_4_ and LXB_4_) as endogenous neuroprotective signals. The canonical role of lipoxins is the regulation of leukocyte and T-cell functions to ensure healthy immune responses[7–10]. However, our recent study established direct protection of RGC and neurons by lipoxins as a new bioaction[6]. In response to retinal excitotoxic stress, formation of LXA_4_ and LXB_4_ in the retina and optic nerve is downregulated, suggesting a homeostatic function. More importantly, inhibition of the lipoxin pathway amplifies RGC death in the retina[6,11] whereas LXB_4_ therapeutic treatment rescued RGC function in a chronic OHT model of neurodegeneration[6]. Notably, LXB_4_, whose mechanism of action is not yet well defined, consistently demonstrated more potent neuroprotective activity than LXA_4_ *in vitro* and *in vivo*[6]. LXB_4_’s action are not mediated by the LXA_4_ receptor, and it has distinct bioactions from LXA_4_ with monocytes and macrophages[12]. The mechanisms through which LXB_4_ exerts its protective effects and the specific cellular targets of LXB_4_ within the retina and optic nerve is not fully understood and was the focus of this study.

A cell type of interest in the cascade of glaucomatous neurodegeneration are microglia, which reside around ganglion cell somata in the retina and their axons in the optic nerve[13–15]. They are in constant communication with astrocytes[16] and continually monitor the retinal and optic nerve environments as part of their essential homeostatic function. Microglial activation and the phenotypic switch from highly ramified, homeostatic microglia to amoeboid-reactive microglia are key features of retinopathy and neurodegeneration[17,18]. Their role in initiating or driving the pathogenesis of glaucoma remains to be clearly defined[19–21]. However, microglia-secreted cytokines regulate astrocyte reactivity in other neurodegenerative disease models including Alzheimer’s disease[22–24].

Using single-cell transcriptomics, bulk RNA-seq, and morphological analysis approaches, we investigated the mechanism and cell targets for LXB_4_ in healthy and OHT-injured retinas. We report that resident microglia in the retina and distal myelinated optic nerve are a cellular target for LXB_4._ After OHT-induced retinal injury, neuroprotective LXB_4_ treatment shifted the functional phenotype of microglia towards homeostasis in the distal myelinated optic nerve and downregulated a key marker of disease-associated microglia. Our findings provide new insights into the functional response of microglia to retinal OHT injury and identify a new cellular target and mechanism for LXB_4_’s neuroprotective actions.

## Methods

### Animals

C57BL/6J and *Alox5^−/−^* (B6.129S2-*Alox5^tm1Fun^*/J, stock number 004155) male mice were obtained from Jackson Laboratory (Bar Harbor, ME). C57BL/6J mice were used as a congenic control for the *Alox5^−/−^* (5-LOX KO) mice. All animal procedures were approved by the Institutional Animal Care and Use Committee (IACUC) at the University of California, Berkeley. Mice were housed in a controlled environment, maintained on a 12-hr light-dark cycle, and provided unrestricted food and water access throughout the study.

### Silicone oil injections

OHT was induced in C57BL/6J and 5-LOX KO mice using an established silicone oil model[25,26]. Briefly, male mice were anesthetized at 8 wks of age using intraperitoneal injection of ketamine/xylazine (100 mg/kg and 10 mg/kg, respectively). A topical anesthetic (0.5% proparacaine hydrochloride; Sandoz, Princeton, NJ) was applied to the eye. Under microscopic guidance, a sterile 31 G paracentesis needle was used to create an incision in the anterior chamber of the eye, ensuring no damage to the iris or lens. The needle was slowly withdrawn to release approximately 1-2 μL of aqueous humor. Subsequently, 1.2 or 1.8 µL of silicone oil (Alcon, Fort Worth, TX) was injected into the anterior chamber using a 33 G Hamilton syringe (Reno, NV). The syringe was slowly withdrawn after holding it for 10 seconds. To minimize silicone oil leakage, the eyelids were gently closed to cover the corneal incision. After the injection, an antibiotic drop (0.3% Tobramycin Ophthalmic Solution, Bausch and Lomb, Laval, Canada) was applied to the eye. Silicone oil was injected into both eyes, and the uninjected eyes from separate groups of animals were used as normotensive controls. Mice were kept on a heating pad until fully recovered from anesthesia. Intraocular pressure (IOP) was measured by Tonometer (TonoLab, Vantaa, Finland) at different time points after dilating the eyes with tropicamide solution (Akorn, Lake Forest, IL) for 10 mins. At designated time points (1, 2, 4, and 6 wks) post-injection, mice were euthanized, and eyes were enucleated in sterile phosphate buffer saline (PBS) at 4 °C. Retina and optic nerve were dissected and stored in 4% paraformaldehyde (4 °C) and TRIzol (−80 °C) for immunostaining and RNA isolation.

### Ocular coherence tomography

Anesthetized mice were maintained on a water-based heating pad at 37 °C. Before imaging, lubricant eye drops were instilled on the eyes (Systane Ultra, Alcon), and pupils were dilated using 0.5% tropicamide (Akorn). A lubricant gel (Tears, Alcon) was used to avoid further drying. A Bioptigen Spectral Domain OCT System (Envisu R2300, Durham, NC) was used for image acquisition. OCT imaging and analysis procedures have been described previously[27,28]. In brief, a rectangular scan with dimensions of 1.8 x 1.8 mm was employed to capture an en-face retinal fundus image centered around the optic nerve head. Each image consisted of 100 B-scan images, with 1536 A-scans for each B-scan. Using a script written in ImageJ software (NIH, Bethesda, MD), masked observers analyzed the retinal layer B-scan images. Retinal layer thickness was quantified from both the left and right locations relative to the center of the optic nerve head. The average value obtained from these locations represented the thickness measurement.

### Electroretinography (ERG)

The mice were subjected to overnight dark adaptation before the ERG measurements. Anesthesia was induced under red light using intraperitoneal injection of ketamine/xylazine (100 mg/kg and 10 mg/kg, respectively). Corneal anesthesia was achieved using topical proparacaine hydrochloride. To dilate pupils, 0.5% tropicamide (Akorn) and 2.5% phenylephrine (Paragon BioTeck, Portland, OR) were applied. ERG measurements were conducted using the Celeris system (Diagnosys LLC, Lowell, MA), employing a range of stimulus intensities from −5.90 to 2.25 log cdm-2. The subdermal needle electrode at the tail served as the ground electrode. The positive scotopic threshold response (pSTR) was elicited using an intensity of −2.50 log cdm-2 to assess RGC function. The pSTR was recorded as the average of 20 repeats with an inter-stimulus interval of 2 seconds. The amplitude at ∼110 ms after stimulus onset was measured and used for analysis.

### LXB_4_ treatment

For scRNA-seq experiments, mice were treated with 1μg of LXB_4_ methyl ester (Cayman Chemicals, Ann Arbor, MI) by intraperitoneal injection (IP) once a day and 1μg of LXB_4_ methyl ester by topical application 3 times a day for 3 days. Vehicle (ethanol) for LXB_4_ methyl ester was removed under a stream of nitrogen, and LXB_4_ methyl ester was resuspended in sterile PBS immediately prior to injection or topical treatment. For RNA-seq experiments, mice were treated with 250 ng of LXB_4_ methyl ester by IP and 25ng of LXB_4_ methyl ester by topical application every other day for 2 wks. For morphometric analysis, mice were treated with 1μg of LXB_4_ methyl ester by IP and 1μg of LXB_4_ methyl ester by topical application once daily for 1 wk. For the sham group, mice were treated with LXB_4_ equivalent volume of sterile PBS via IP and a topical route. LXB_4_ treatment was initiated ∼15 mins before OHT induction.

### Quantitative PCR

Total RNA was isolated from retinas using TRIzol extraction method (Invitrogen, Waltham, MA). mRNA was converted to cDNA using an iScript cDNA synthesis kit (Bio-Rad, Hercules, CA). Transcripts for *C5ar1, Clec4a2, C3ar1, Ccl5, Tnf-α, Cxcl10,* and *Cd68* were quantified by using GoTaq PCR master mix (Promega, Madison, WI) in OneStep Plus qPCR (Applied Biosystems, Waltham, MA) system by 2^−ΔΔCT^ method.

### Immunostaining

For whole mounts of the retina and optic nerve, the tissues were dissected and fixed in 4% PFA at 4 °C overnight. The next day, tissues were blocked and permeabilized in blocking buffer (10% normal donkey serum + 2% Triton x-100) for 24 hrs at 4 °C. Further, tissues were incubated with primary antibodies at 1:1000 dilution (anti-Iba1, Cell Signaling (Danvers, MA); anti-CD68, BioLegend (San Diego, CA); anti-RBPMS, Phosphosolutions (Aurora, CO)) for 72 hrs at 4 °C. Tissues were rinsed three times with washing buffer (PBS+ 0.25% Triton x-100) solution, each for 10 mins, on a rocker at room temperature. Secondary antibodies were diluted in blocking buffer at 1:2000 dilution (Alexa Fluor 594 and Alexa Fluor 488, Invitrogen), and tissues were incubated with a secondary antibody cocktail overnight at 4 °C. The next day, whole mounts were rinsed three times in a washing buffer and incubated with DAPI (1:5000, Invitrogen) for 10 mins at room temperature. Whole mounts were mounted using gold antifade mounting medium (Invitrogen) on a slide in a coverslip grove. For staining of optic nerve sections, the optic nerves were dissected and fixed in 4% PFA at 4 °C overnight. The next day, optic nerves were washed in PBS and dehydrated in a sucrose gradient (10%, 20%, and 30% sucrose) before embedding them in optimal cutting temperature medium (Thermo Fisher, Waltham, MA). 10µm sections were taken in Leica CM1900 cryostat (Wetzlar, Germany). Sections were washed in PBS, blocked, and permeabilized in blocking buffer (10% normal donkey serum + 0.25% Triton x-100) for 1 hr. Sections were then incubated in primary antibodies (anti-CD74, 1:100, BioLegend; anti-Iba1, 1:100; anti-p-PI3K, 1:100, Invitrogen) dissolved in blocking buffer overnight at 4 °C. The next day, sections were washed in PBS three times for 10 mins each and were incubated with secondary antibodies (Alexa Fluor 488 and Alexa Fluor 594, 1:200) for 2 hrs at room temperature. Sections were washed with PBS, incubated with DAPI (1:2500) for 10 mins, and mounted using FluorSave™ mounting media (Sigma Aldrich, St. Louis, MO).

### Image Analysis

RGCs were counted from whole mounts using a macro written in ImageJ, which sets the auto threshold, performs water-shedding and counts RGCs by Analyze Particles(). Two images were acquired from each flank of the retinal whole mount (Supplementary Figure 2A), and the mean of the 8 image counts represents the count for one retina. After adjusting the threshold, Iba1 and CD74 positive microglia were counted from optic nerve sections using Analyze Particles(). For CD74 and phospho-PI3K co-localization, single cells were cropped from whole optic nerves sections and were colocalized using Coloc2 and BIOP plugins in ImageJ. Pixel intensities were measured using the Measure() function in ImageJ software. Pearson’s correlation coefficient was calculated in RStudio by cor.test().

### Confocal microscopy

Images of microglia whole mounts and sections were acquired using a Zeiss LSM710, Axio Imager 2 with Plan-Apochromat 20x objective, 0.8 NA and Plan-Apochromat 63x objective, 1.4 NA. Z-stacks of the retina and optic nerve whole mounts and sections were taken at 1024×1024 resolution. For retina, at least 4 different images, one from each retinal whole mount flank (Supplementary Figure 2A) were taken, and for the optic nerve, at least 3 different images (Supplementary Figure 2B) were taken.

### Microglia feature analysis

The features of microglia morphology were analyzed by the method published by Heindl et al[29] using custom scripts in MATLAB (R2022a, MathWorks, Natick, MA), which relies on the Image Processing Toolbox and Statistics and Machine Learning Toolbox for its functionalities. The fully automated analysis of morphological features extracted from confocal image stacks of Iba1 stained microglia involved four primary steps. First, the image quality and preprocessing were controlled to ensure reliable results. Second, microglial cells were segmented from the background, and within each cell, further segmentation was performed to distinguish the nucleus, soma, and branches. Third, a skeleton representation is constructed to capture the spatial structure of the cell bodies and branches. Finally, morphological features were extracted using the properties derived from the cell surface area, volume, and skeleton. The output feature file was imported into RStudio v.4.2.0, and the mean values for individual retina and optic nerves were calculated for each replicate. Mean values were plotted using GraphPad Prism 9 software (La Jolla, CA).

### MorphOMICs analysis

#### Reconstruction of microglia morphology

Microglia morphology was reconstructed using Imaris 9.2.v (Oxford Instruments, Abingdon, UK) software. Briefly, raw z-stack confocal image files were imported into Imaris, and the surface module was used to construct the surfaces on the Iba1 channel by setting smoothing=1. The new masked channel was created on the surface. Further, microglial processes were analyzed in three dimensions using a filament-tracing plugin. Starting points (soma) for tracing were identified by setting a maximum diameter of 12 µm, and the seeding points (dendrites) were identified by setting a diameter of 1 µm. Following the tracing process, we manually excluded cells located at the image border that were only partially traced, thereby ensuring that these cells were not included in the analysis. However, it is highly challenging to manually remove minor artifact filament generated by Imaris. The .ims files were then converted to .swc files for individual microglia using the ImarisReader toolbox (https://github.com/rcubero/Matlab_Imaris_converter). To count microglia, soma statistics were used after using filament tracer plugin.

#### MorphOMICs pipeline

Microglia morphology was mapped using MorphOMICs (https://github.com/siegert-lab/MorphOMICs). MorphOMICs uses microglia’s topological morphology descriptor (TMD) combined with bootstrapping and dimension reduction techniques, Principal Component Analysis (PCA) and Uniform Manifold Approximation and Projection (UMAP), to visualize unsupervised clustering of microglia from different treatment conditions. The bootstrap sample size was 100, and the number of bootstraps collected was 400. The first ten principal components were used as input to UMAP with n_neighbors□=□50, min_dist□=□1.0, and spread□=□3.0. Importantly, bootstrapping and PCA reduce the impact of minor artifact filaments introduced by Imaris in the final clustering outcome.

#### Monocle trajectory analysis

Each bootstrapped sample is represented by an array of 10,000 pixels from its persistence image, therefore a pixel from a sample can be seen as analogous to a gene from a cell within a single-cell transcriptomic setting. Pixels in proximity covary with one another in a manner similar to how genes may covary.

A pseudo-temporal trajectory-inference algorithm called Monocle[30–32] was used. Monocle uses a partitioned approximate graph abstraction-like algorithm for Louvain clustering, which learns a principal graph using reversed graph embedding to generate lineages and pseudotimes (https://github.com/cole-trapnell-lab/monocle3/).

The bootstrapped samples from MorphOMICs were used as input for Monocle. The principal components were obtained using preprocess_cds with num_dim = 10, after which UMAP dimension reduction was performed using reduce_dimension with umap.metric = ‘manhattan’, umap.min_dist = 1.0, and umap.n_neighbors = 50. Clusters were determined using cluster_cells with cluster_method□=□‘leiden’, and the pseudo-temporal trajectory was obtained using learn_graph with use_partition□=□FALSE and close_loop□=□FALSE.

### Single-cell transcriptomics

#### Single-cell dissociation

Retinas were dissociated into single-cell suspensions using a papain dissociation kit (Worthington, Columbus, OH). Briefly, dissected retinas were placed in papain solution for 30 mins at 37 ° C with exposure to 5% CO_2_ in the incubator. Retinas were gently tapped and incubated for 15 more mins for complete dissociation. The reaction was quenched by adding ovomucoid inhibitor solution, and cells were collected after centrifugation and dissolved in resuspension buffer containing PBS+5%BSA+ DNase to form a single-cell solution of cells. Cells were passed through a 40μM cell strainer (Sigma) to remove debris and clumps. Before processing the cells for single-cell transcriptomics, rod cells were depleted to enrich the rest of the population. Cells were incubated with Biotin-CD133 antibody for rod-specific labeling and were depleted in the magnetic column after anti-biotin-magnetic bead labeling (Miltenyi Biotech, Bergisch Gladbach, Germany).

#### Barcoding and Library Preparation

Barcoding and library preparation was achieved using a 10X Chromium Single Cell 3’ reagent (v3.1 chemistry, 10x Genomics, Pleasanton, CA), according to the manufacturer’s instructions. Briefly, cells were loaded on the chromium chip to create gel beads containing unique oligo barcodes with single cells, creating cell-gel droplets. RNA from each cell was captured within the droplets, and cell-specific barcodes were linked to their respective transcripts. Reverse transcription and amplification of captured RNA were performed to generate cDNA libraries. Quality of cDNA preparation was checked using Bioanalyzer (Applied Bioscience)

#### Sequencing

The cDNA libraries were subjected to high-throughput sequencing on the Illumina Novaseq S1 100SR platform (San Diego, CA). Sequencing reads containing cell-specific barcodes and transcript information were obtained for each cell. Raw files were demultiplexed using Illumina bcl2fastq2 software for downstream analysis. Library preparation and scRNA-seq were performed at the QB3 Genomics Core Facility, UC Berkeley, Berkeley, CA, RRID: SCR_022170.

#### Downstream Data Analysis

The sequencing reads were processed using Cellranger software (10x Genomics). Cellranger ‘count’ pipeline aligned the sequencing reads to the existing mouse genome (mm10, University of California Santa Cruz). Each read was assigned to its respective cell barcode, enabling cell identification and quantification. The output matrix was used for further downstream analysis and clustering of the cell populations in RStudio v.4.2.0 using Seurat V4.0[33]. Briefly, quality control and filtering were performed after determining the percentage of mitochondrial transcripts. The data were normalized using the NormalizeData() function, ensuring that gene expression values were comparable across cells. Variable features were selected using the FindVariableFeatures() function to identify the genes that exhibited significant variation and contributed to heterogeneity within the dataset.

The FindIntegrationAnchors() function was executed on both objects to enable comparisons across different datasets. This process identified cell-to-cell pairings between the two datasets based on the first 30 principal components (PCs). The objects were then integrated using the IntegrateData() function on the anchorset and the first 30 PCs. This integration step combined the gene expression profiles from different datasets while preserving the underlying biological variability.

After integration, counts were scaled using the ScaleData() function to normalize the expression values across cells. Principal components (PCs) were computed using the RunPCA() function, capturing the major sources of variation within the integrated dataset. Furthermore, clustering analysis was performed by executing the FindNeighbors() function on the first 30PCs, identifying the nearest neighbors for each cell based on their gene expression profiles. Subsequently, the FindClusters() function (resolution-0.8) was used to assign cells to distinct clusters based on their similarity in gene expression. A uniform manifold approximation and projection (UMAP) was constructed using the RunUMAP() function on the first 30 PCs, allowing for the visualization of the cells in a lower-dimensional space while preserving the global structure of the data.

To assign cell types to the identified clusters, markers specific to each cluster were identified using the FindAllMarkers() function and compared to known cell-type markers[34–36], facilitating the annotation of the clusters with specific cell types. Differential gene expression in microglial cells was determined using the FindMarkers() function. Data were visualized using VlnPlot() and DotPlot() functions.

### Bulk RNA Sequencing

Total RNA was isolated from optic nerves by TRIzol method (Invitrogen) and RNA quality was measured on Bioanalyzer (Applied Bioscience). mRNA was converted to cDNA using SMARTer v4 Ultra Low Input RNA Kit (Clontech, Mountain View, CA). A Diagenode Bioruptor Pico was used to fragment the cDNA, and libraries were generated using the KAPA Hyper Prep Kit for DNA (Roche, Basel, Switzerland) for sequencing on a NOVAseq S4 flow cell (Illumina). Library preparation and RNA-seq were performed at QB3 Genomics, UC Berkeley, Berkeley, CA, RRID: SCR_022170. The raw sequencing reads were demultiplexed by Illumina bcl2fastq2 software, and read quality was assessed using FastQC v.0.11.9. Adapters were subsequently trimmed from the reads using Trim Galore v.0.6.6. The processed reads were then aligned to the mouse genome (mm39, University of California Santa Cruz) using the STAR alignment tool v.2.7.1a and read counts for each gene were obtained using FeatureCount v.1.5.3. The resulting feature count matrix was imported into RStudio v.4.2.0, and normalization and differential gene expression analyses were conducted using the DeSeq2 package. Heatmap and venn diagram was generated using pheatmap() and ggvenn() packages in RStudio v.4.2.0.

Additionally, pathway enrichment analysis was performed using the clusterProfiler package, allowing for the identification of biological pathways enriched with differentially expressed genes. The interaction network of genes was created by String DB, and the Highly interacting network of genes was analyzed by MCODE v. 2.0.2 in Cytoscape v.3.9.1 software. Pathway enrichment of the MCODE-generated networks was performed using ClueGO v. 2.5.9 in Cytoscape v.3.9.1, using a significance cutoff of p < 0.05.

### Statistical analysis

Student’s t-test was used to determine the significance of differences (p<0.05) between the two groups. One-way analysis of variance with post-hoc Tukey’s multiple comparison tests was used to compare multiple groups. Values are presented as mean ± SEM (standard error of the mean).

## Results

### LXB_4_ regulates microglia functional responses in the healthy and glaucomatous retina

Our previous study established endogenous formation of LXB_4_ in the healthy retina and its neuroprotective effect in a rat suture model of OHT by directly acting on RGCs[6]. Data on the mechanism for LXB_4_’s bioactions and cellular targets are sparse. Hence, we treated healthy mice with LXB_4_, *in vivo*, to investigate its homeostatic actions and uncover cellular targets. Cell-specific actions were defined by single-cell transcriptomics (scRNA-seq) analysis (Figure 1). After unsupervised clustering, supervised cell type identification, and differential gene expression analysis (|Log2FC|>1, p.adjust<0.05), identified two cell types highly regulated by LXB_4_ treatment in the healthy retina (Figure 1A), namely microglia (38 differentially expressed genes) and astrocytes (36 differentially expressed genes). We recently reported that LXB_4_ acts directly on astrocytes and regulates their function [37], hence regulation of microglia in healthy retina by LXB_4_ was unexpected and the focus of this study. We analyzed differentially expressed genes in microglia (Figure 1B) with pathway enrichment analysis focused on downregulated genes (Log2FC< −1, p.adjust<0.05, total 27 genes). The analysis identified downregulation of genes for antigen processing and presentation (Figure 1C), which indicates that LXB_4_ treatments downregulates microglia pathways that have been linked to neuroinflammation. More importantly, some of highly downregulated genes identified in enrichment analysis were *C5ar1*, *Clea4a2*, *Entpd1*, *Il6ra*, and *CD37* (Figure 1D) which have been identified as part of the “sensome” encoding proteins expressed on ramified processes of microglia[38] were primarily expressed in retinal microglia (Figure 1E). The function of the sensome is regulation of microglia homeostatic and reactive functions in response to microenvironmental cues[18]. The five genes downregulated by LXB_4_ are linked to the transition from a homeostatic phenotype to a reactive microglia phenotype[18]. The role of microglia in the cascade leading to RGC degeneration is of great interest[39] and remains to be defined.

**Figure 1:**
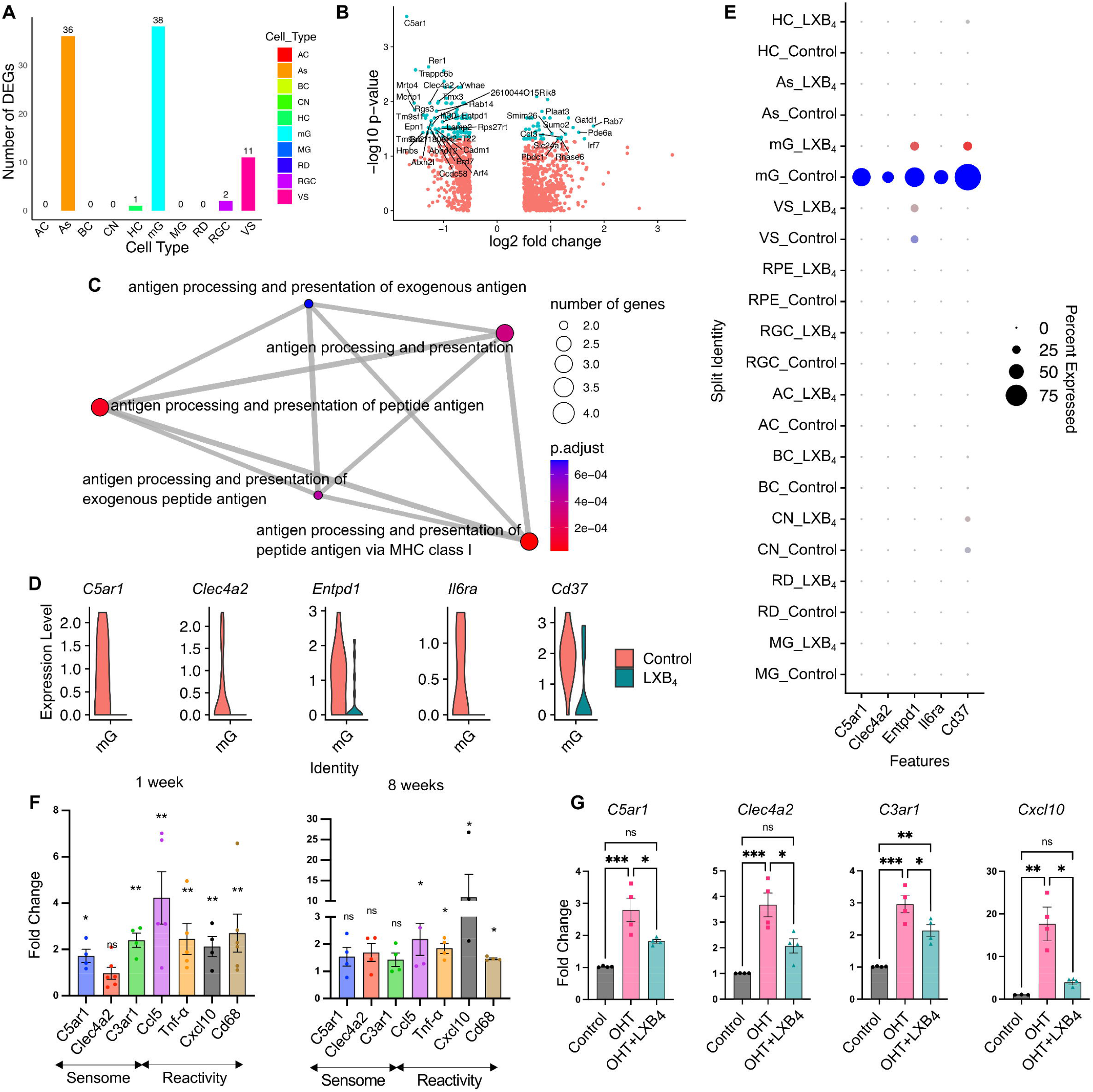
LXB_4_ targets retinal microglia in healthy and moderate ocular hypertension conditions. **(A)** Number of differentially expressed genes with |Log_2_ Fold Change (FC)|> 1 and p.adjust<0.05 after LXB_4_ treatment are plotted across the retinal cell types. AC: amacrine cells, As: astrocytes, BC: bipolar cells, CN: cones, HC: horizontal cells, mG: microglia, MG: muller glia, RD: rods, RGC: retinal ganglion cells, VS: vascular cells. **(B)** Volcano plot of differentially expressed genes in microglia comparing LXB_4_ treatment versus healthy condition. Genes with |Log_2_ FC|> 0.5 change in expression are plotted. Significant (p.adjust< 0.05) changes in cyan and not significant in red. Genes with |Log_2_FC|>1 significant changes are labeled. (**C)** Gene ontology pathway enrichment map for genes downregulated (Log_2_FC < −1, p.adjust <0.05) by LXB_4_ in the healthy retina. (**D)** Violin plot of selective sensome genes downregulated in microglia (mG) by LXB_4_ treatment in healthy retinas. (**E)** Dot-plot of sensome genes expressed in retinal cells of healthy controls (blue) and after LXB_4_-treatment (red). Size of dot represents percentage of cells expressing the gene and color intensity represent average expression. (**F)** Quantitative PCR analysis of sensome and microglia reactivity gene expression from whole retina after 1 and 8 wks of moderate OHT. Significance was determined by unpaired t-test (*p< 0.05; **p<0.01; ns, not significant, n=4); each replicate was pooled from two retinas. (**G)** Quantitative PCR analysis of microglia sensome and inflammatory gene expression from whole retina with or without LXB_4_ treatment for 3 wks during OHT compared to normotensive retinas. Data were analyzed by One-way ANOVA with Tukey’s multiple comparison test (*p< 0.05; **p<0.01; ***p<0.001; ns-, not significant, n=4). Each replicate is 2 retinas pooled. Data is presented as mean ± SEM.

To investigate LXB_4_ regulation of microglia function during OHT-induced RGC degeneration, we used the established silicone oil model of chronic moderate OHT[25,40,41] (Supplementary Figure 1A-C) and analyzed microglial sensome and reactivity genes by qPCR. Importantly, a signature sensome gene (*C5ar1*) that was downregulated by LXB_4_ in the healthy retina (Figure 1D) was highly upregulated (171%, p=0.0190) under chronic OHT at the 1 wk time point, along with markers of microglial reactivity genes (*Ccl5, Tnf-*_α_*, Cxcl10, Cd68*) after 1 or 8 wks (∼200 to 1000%) (Figure 1F). Further, sensome genes (*C3ar1*, 295%, p= 0.0001; *C5ar1*, 279%, p= 0.0007; and *Clec4a2*, 367%, p= 0.0005) were highly upregulated after 4 wks of OHT. Importantly, LXB_4_ treatment significantly downregulated expression of sensome genes (*C3ar1*, 27.7%, p= 0.0305; *C5ar1*, 35.1%, p= 0.0271; and *Clec4a2*, 43.5%, p= 0.0140) and the inflammatory marker *Cxcl10* (77.8%, p= 0.0132) in the retina (Figure 1G). It is important to note that various components of the microglia sensome (*C3ar1, C5ar1, Clec4a2*) may exhibit diverse and temporarily defined activation in responses to sustained OHT stress. The role and function of sensome genes, including *Clec4a2*, in OHT is currently unknown. These data identify and suggest microglia as a target for LXB_4_ regulation in the healthy normotensive retina and during moderate OHT stress through direct or potentially indirect mechanisms.

### Moderate OHT regulates microglia phenotype in both the retina and optic nerve

Since we identified microglia as a new cell target for LXB_4,_ and the roles of microglia in glaucomatous conditions are complex[19–21], we aimed to better understand the dynamic regulation of microglia function during moderate OHT. While transcriptomics[42] and proteomics[43] studies advanced our understanding of signaling mechanisms in microglia, morphology is a key functional characteristic of microglia behavior and phenotype[44–46], reflecting their spectrum of biological functions[46]. New analytical methods to quantify microglial morphology in detail [29,47] have made it feasible to comprehensively measure the morphological features of microglia and relate them to distinct phenotypes[29,47]. We analyzed the effects of moderate OHT on microglia functional responses in the retina and optic nerve using the methods established by Heindl et al. (2018)[29] and Colombo et al. 2022[47]. The morphologies of retinal and optic nerve microglia stained with Iba1 were analyzed after 2, 4, and 6 wks of OHT. First, we used a feature extraction tool from Heindl et al. [29], which extracts predefined set of morphological features (Sphericity, Circularity, Volume of nodes, Total number of nodes, etc.) from Iba1 stained microglia image stack. In the retina, out of 14 features of microglia morphology analyzed, ∼14% of features (Reduction in length of branches and Betweenness of nodes) were significantly changed (p<0.05) as early as 2 wks when compared to normotensive control retinas (Supplementary Figure 2C). With sustained OHT, 78% of features were significantly changed (p<0.05) at the 4 wks timepoint. At 6 wks, ∼93% of features were changed (p<0.05), representing a time-dependent change in the phenotypes of retinal microglia (Supplementary Figure 2C). Next, we analyzed whole mount Iba1-stained microglia in the myelinated optic nerve. None of the optic nerve microglial features significantly changed at the early 2 wks point (Supplementary Figure 3). However, at 4 wks, ∼64% of analyzed features exhibited statistically significant changes (p<0.05) when compared to the normotensive controls. Furthermore, when comparing the 4 wks data to the 2 wks time point, around 21% of the features showed statistically significant differences (p<0.05), establishing time-course specific increase in microglia morphology changes (Supplementary Figure 3). These results indicate that moderate OHT in both the retina and distal myelinated optic nerve led to specific and dynamic changes in microglia phenotypes.

Next, the morphOMICs method developed by Colombo et al. (2022)[47] was used to capture the complex variability in microglia morphology and to define the distinct microglia subpopulations present in the retina and optic nerve. MorphOMICs involves the conversion of a 3D microglia structure to a 2D persistence barcode[48] and applying bootstrapping and dimension reduction methods to visualize the distinct clusters of microglia with morphological differences[47]. The persistence barcode is generated using Topological Morphology Descriptor (TMD)[48], which maps the start and end of microglia processes as their distance from the soma, enabling it to be a more sensitive, Omic and efficient analysis than the feature extraction tool[29]. After analyzing the retinal microglial whole mount images at different time points of moderate OHT, we observed that retinal microglia clustered differentially at different time points, denoting unique phenotypes at different stages of moderate OHT (Figure 2A-C). These results were concurrent with the feature extraction method, which revealed unique time-course specific morphological characteristics of microglia. To establish the time-dependent changes in microglia morphology, we used pseudo-temporal trajectory-inference algorithm (Monocle3) analysis[30–32], which utilizes reversed graph embedding to generate lineages and pseudotime trajectories. Normotensive control microglia represent the starting point of the trajectory. At 4 wks of OHT, microglia in the retina were farthest from the normotensive control microglia, denoting the homeostatic-to-reactive phenotype spectrum (Figure 2D). Retinal microglia at the 6 wks time point of OHT aligned between the 2 wks and 4 wks time points on the trajectory (Figure 2D), suggesting a gradual reversal of a microglia inflammatory functional phenotype in the retina. It is important to note that there were no significant changes in total number of microglia in the retina at these time points (Supplementary Figure 4A). In addition, we used the microglia reactivity marker CD68[49] to identify reactive microglia at different stages of OHT. Retinal microglia expressed CD68 protein as early as 2 wks and the reactivity marker was expressed throughout the entire time course of 6 wks, with maximum protein expression at 4 wks in the retinal microglia population (Figure 2E, F).

**Figure 2:**
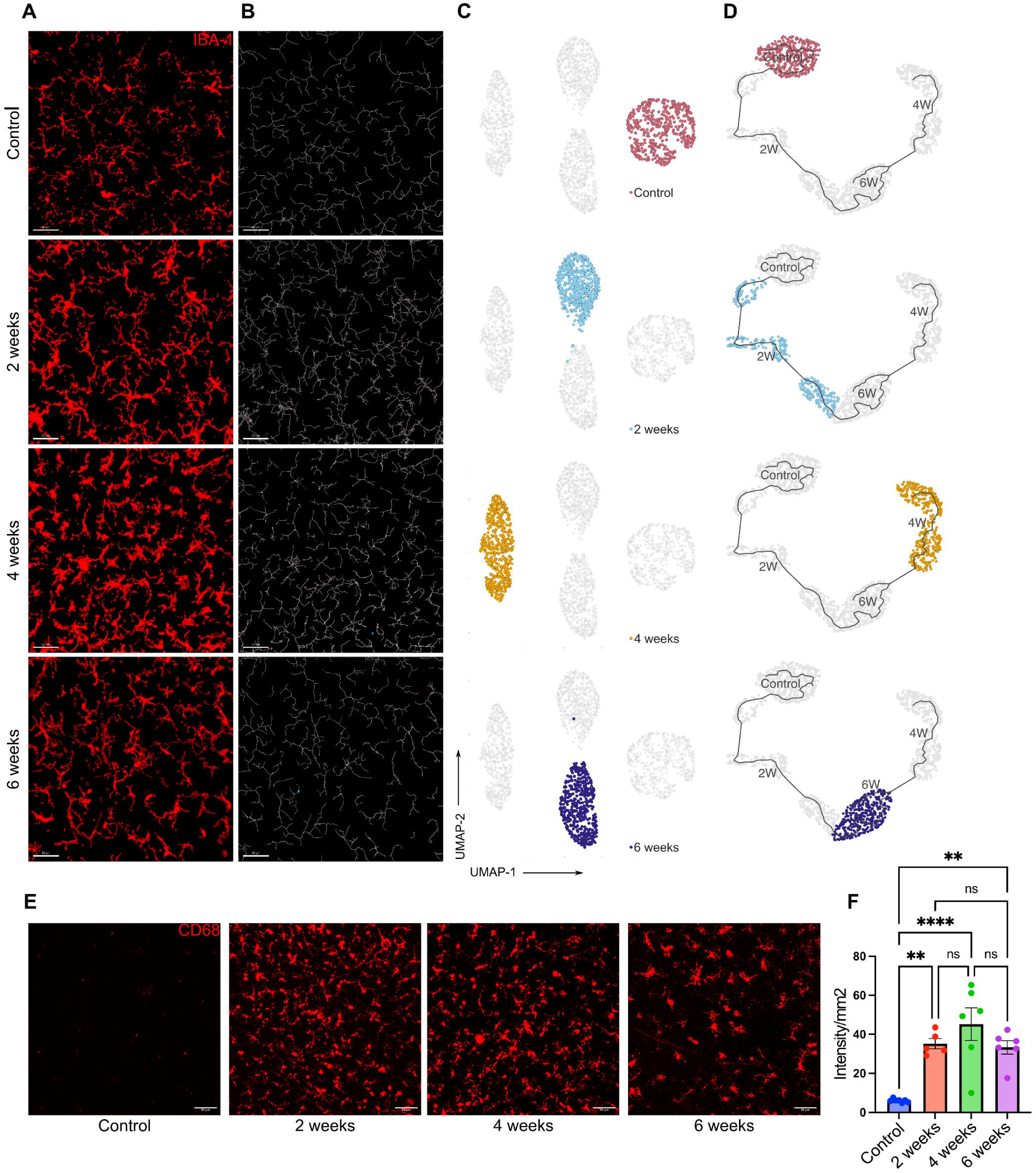
Dynamic changes in retinal microglia morphology during moderate OHT. **(A)** Representative confocal micrograph of Iba1 (red) stained retinal microglia at different time points of moderate OHT (scale bar-50μm). (**B)** Representative filament images of microglia morphology over the time course of OHT, generated from the respective Iba1-stained images using IMARIS (scale bar-50μm). (**C)** UMAP plot displaying the clustering of microglia populations at different time points of OHT. Each dot depicts 100 microglia bootstrapped for their topological morphology descriptor, and 400 dots per group. (**D)** UMAP plot of pseudotime trajectory of microglia population morphology during time course of moderate ocular hypertension; each time point is represented by a different color. (**E)** Representative confocal micrographs of CD68-stained retinal microglia at different time points of OHT (scale bar-50μm). (**F)** Quantification of CD68 expression in retinal whole mounts in normotensive controls (n=6) and 2 wks (n=5), 4 wks (n=6), and 6 wks (n=6) of moderate OHT. Data were analyzed by One-way ANOVA with Tukey’s multiple comparison test (**p<0.01; ****p<0.0001; ns, not significant). Each dot depicts the mean CD68 expression from a single retina. Data presented as mean ± SEM.

Next, we performed morphOMICs analysis of optic nerve microglia during the time course of OHT. In contrast to retinal microglia, normotensive control and 6 wks optic nerve microglia populations clustered together (Figure 3A-C), while microglia populations from the 2 and 4 wks timepoints clustered separately from both the normotensive control cluster and the 6 wks cluster (Figure 3C). To further understand these time-dependent phenotypic changes in optic nerve microglia, we performed pseudotime trajectory analysis. Similar to the retina, the microglia populations at 4 wks were farthest on the trajectory compared to the microglia population from healthy optic nerves (Figure 3D), establishing the 4 wks timepoint as the maximal microglia functional response to OHT in both the remote optic nerve and retina. More importantly, similar to the retina, microglia populations at the 6-wks time point partially clustered near the normotensive control cluster in the pseudotime trajectory analysis (Figure 3D). Similar to the results from feature extraction, optic nerve microglia showed homeostatic-like morphology at 6 wks of moderate OHT (Figure 3D), denoting similar regulation of microglia functional phenotypes at two distinct sites during the time course of moderate OHT. There was no change in the total number of microglia (Supplementary Figure 4B) in the optic nerve and the reactivity marker CD68 was upregulated over the entire time course of OHT at the remote myelinated optic nerve site (Figure 3E, F). Collectively, these data indicate the similar regulation of retinal and optic nerve functional microglia phenotypes under moderate OHT.

**Figure 3:**
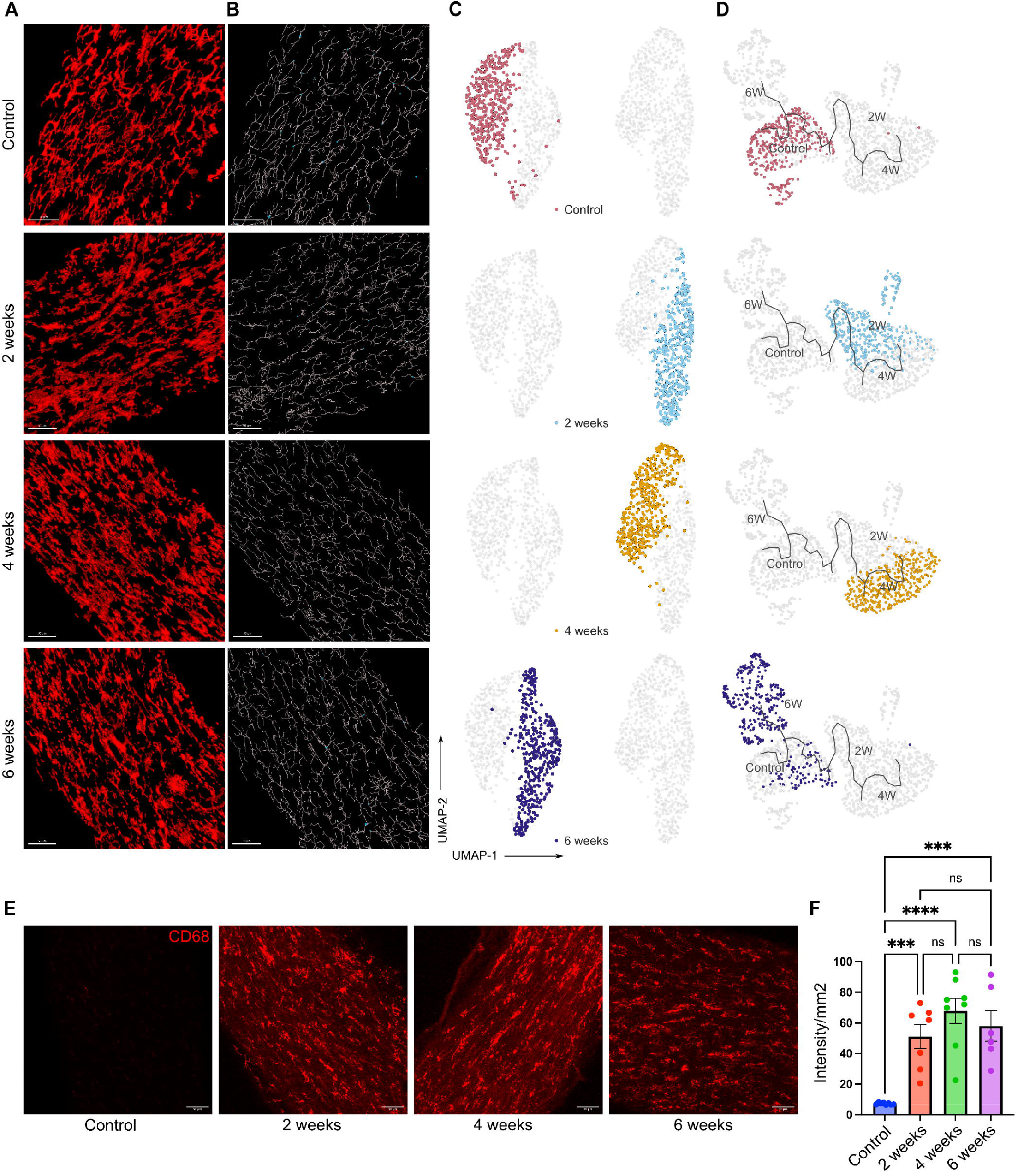
Dynamic changes in optic nerve microglia phenotypes during moderate OHT. **(A)** Representative confocal micrograph of Iba1 (red) stained optic nerve microglia at different time points of OHT (scale bar-50μm). (**B)** Representative filament images of microglia morphology during the time course of OHT, generated from the respective Iba1-stained images using IMARIS (scale bar-50μm). (**C)** UMAP plot displaying clustering of optic nerve microglia during time course of OHT. Each dot depicts 100 microglia bootstrapped for their topological morphology descriptor, and 400 dots are shown per group. (**D)** UMAP plot of pseudotime trajectory of microglia morphology for the time course of moderate ocular hypertension for each time point(**E)** Representative confocal micrographs of CD68-stained optic nerve microglia at different timepoints of OHT (scale bar-50μm). (**F)** Immunohistochemical quantification of CD68 expression in optic nerve whole-mounts for normotensive controls (n=6), 2 wks (n=7), 4 wks (n=8), and 6 wks (n=6) of OHT. Data were analyzed by One-way ANOVA with Tukey’s multiple comparison test (***p<0.001; ****p<0.0001; ns, not significant). Each dot depicts the mean CD68 expression from a single retina. Data presented as mean ± SEM.

### Neuroprotective LXB_4_ regulates optic nerve microglia during retinal OHT insult

Both retinal and optic nerve microglia transition from homeostasis to distinct states and dynamic reactivity phenotypes in response to sustained OHT. scRNA-seq data suggest that LXB_4_ regulates the microglia homeostatic phenotype in the healthy retina. We used a silicone oil volume that induces severe OHT retinal injury (40% peripheral RGC loss) by 2 wks (Supplementary Figure 1D-H) to investigate the actions of LXB_4_ microglia function in response to significant RGC injury. LXB_4_ treatment significantly reduced loss of the nerve fiber layer after 1 wk (∼55% improvement from OHT group) based on ocular coherence tomography (OCT) measurements (Figure 4A, B). Microglia morphology in the retina and optic nerve was analyzed by feature extraction tool and morphOMICs as described above, to define LXB_4_ regulation of microglia function. Analysis of retinal microglia following severe OHT (1 wk) by feature extraction tool from Heindl et al. [29] revealed significant changes (p<0.05) in ∼93% of the analyzed features. LXB_4_ treatment during severe OHT did not significantly affect the morphology of retinal microglia (Supplementary Figure 5). In sharp contrast, LXB_4_ treatment induced or maintained a normotensive morphological phenotype in approximately 57% of features (out of the 7 analyzed, #p<0.05) at the distal myelinated optic nerve that were altered by OHT (Supplementary Figure 6).

**Figure 4:**
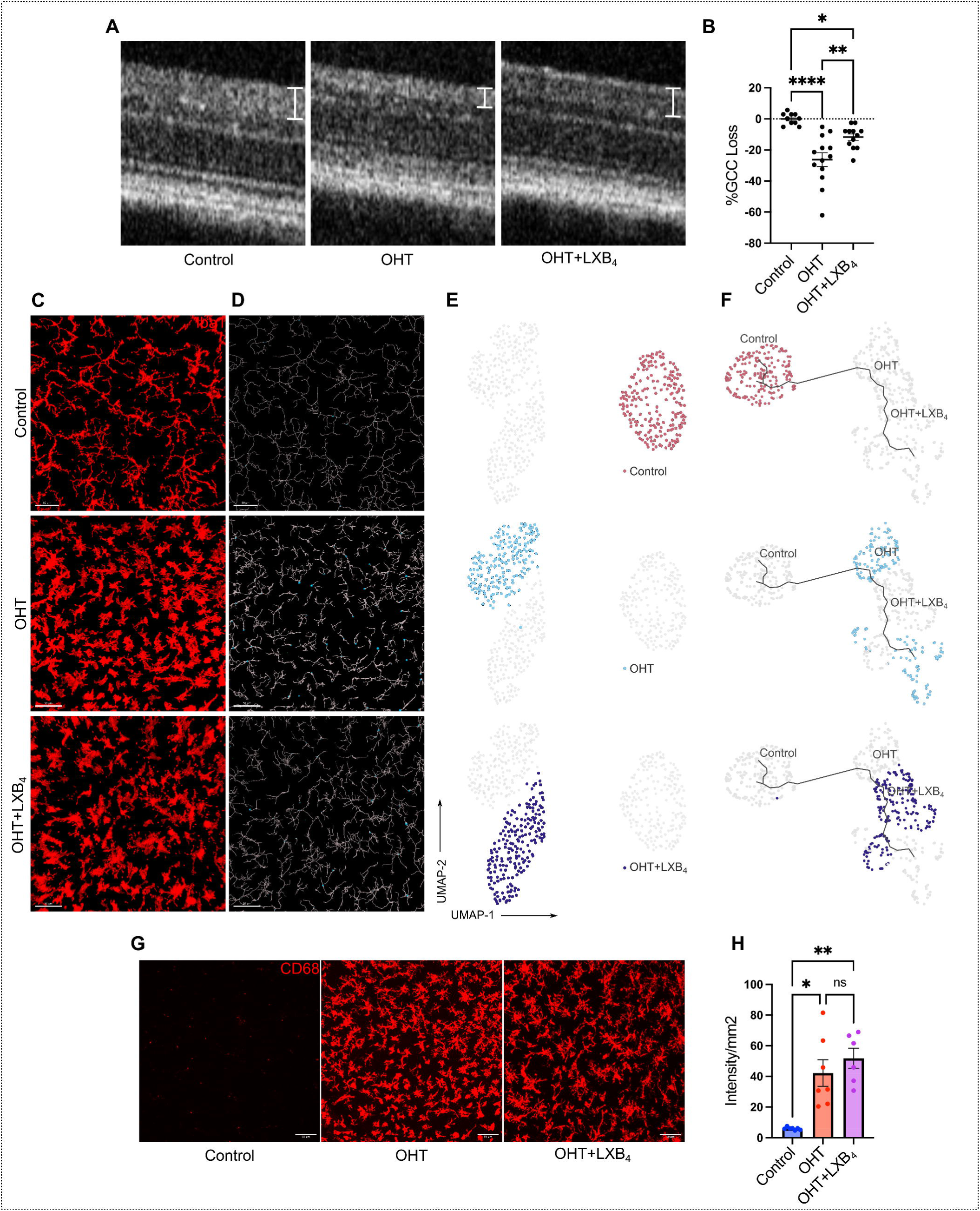
Neuroprotective LXB_4_ treatment did not affect retinal microglia phenotype during severe OHT. **(A)** Representative OCT images of retinal layers for a normotensive control, OHT, and OHT+LXB_4_. Ganglion cell complex (GCC) thickness is marked by line segments. **(B)** Scatter plot quantification of GCC layer thickness for normotensive controls (n=10), OHT (n=13), and OHT+LXB_4_ (n=12). Data were analyzed by One-way ANOVA with Tukey’s multiple comparison test (*p<0.05; **p<0.01; ****p<0.0001). Each dot depicts the percentage of GCC loss. (**C)** Representative confocal micrograph of Iba1 (red) stained retinal microglia in normotensive control, OHT, and OHT+LXB_4_ (scale bar-50μm). (**D)** Representative filament images of microglia morphology generated from the respective Iba1-stained images using IMARIS (scale bar-50μm). (**E)** UMAP plot displaying clustering of optic nerve microglia in normotensive, OHT, and OHT+LXB_4_ groups. Each dot depicts 100 microglia bootstrapped for their topological morphology descriptor, and 200 dots per group. (**F)** UMAP plot of pseudotime trajectory of microglia morphology; all groups are displayed at once. (**G)** Representative confocal micrographs of CD68-stained retinal microglia for normotensive controls, OHT, and OHT+LXB_4_ (scale bar-50μm). (**H)** Quantification of CD68 expression in retinal whole mounts for normotensive control (n=6), OHT (n=6), and OHT+LXB_4_ (n=6). Data were analyzed by One-way ANOVA with Tukey’s multiple comparison test (*p<0.05; **p<0.01; ns, not significant). Each dot depicts the mean CD68 expression from a single retina. The experiment was concluded at 1wk of severe OHT and LXB_4_ treatment. Data presented as mean ± SEM.

MorphOMICs was used to analyze the complex and dynamic changes in distinct microglia populations in detail in the retina and distal myelinated optic nerve during severe OHT and LXB_4_ treatment. In the retina with severe OHT, microglia were uniquely clustered compared to the microglia population from normotensive retinas (Figure 4C-E). MorphOMICs analysis (Figure 4C-E) and pseudo-time trajectory (Figure 4F) established that LXB_4_ treatment during acute and severe OHT did not result in significant morphological changes in retinal microglia populations as they clustered together in the analyses. Severe OHT with or without LXB_4_ treatment also demonstrated a significant increase in the total number of microglia in the retina at 1 wk compared to the normotensive control retina (Supplementary Figure 4C). In addition, LXB_4_ treatment did not reduce CD68 staining, a marker of microglia reactivity in the retina (Figure 4G, H). Taken together, these results indicate that neuroprotection by LXB_4_ treatment during acute and severe OHT is not associated with regulating the retinal microglia phenotype.

Next, we analyzed the potential effect of LXB_4_ treatment on microglia phenotype in the distal myelinated optic nerve. MorphOMICs identified unique clusters of microglia populations in all three groups, namely normotensive, OHT, and OHT with LXB_4_ treatment. Optic nerve microglia from normotensive retina were a single and unique cluster (Figure 5C), while optic nerve microglia populations from severe OHT were spatially distinct and separated into two clusters (Figure 5C). These findings establish that severe OHT induces distinct and rapid morphological changes in microglia populations in the distal optic nerve. Interestingly, unlike in the retina, morphOMIC analysis revealed that LXB_4_ treatment induced a unique single microglia population in the optic nerve, which was a distinct cluster from the sham-treated mice with severe OHT (Figure 5C). Trajectory analysis confirmed that the spatially unique subpopulation of optic nerve microglia in mice with severe OHT were the farthest morphological distance (Figure 5D) from the microglia population in normotensive control eyes. The unique microglia population that was induced by LXB_4_ treatment was closest to the normotensive microglia population in pseudotime trajectory analysis (Figure 5D), i.e., morphological towards the homeostatic phenotype. Unlike in the retina, the total number of microglia in the optic nerve did not change in response to severe OHT (Supplementary Figure 4D). Next, we analyzed microglial reactivity in the optic nerve using CD68 as a reactivity marker. Immunohistochemistry (IHC) analysis demonstrated that severe OHT induced a high expression (297% increase, p=0.0007) of CD68 in microglia in the distal myelinated optic nerve (Figure 5E, F) compared to optic nerves from normotensive eyes. More importantly, LXB_4_ treatment significantly reduced (88%, p=0.0343) CD68 expression after 1 wk of severe OHT (Figure 5E, F). These data are consistent with the morphOMIC analysis, which demonstrates that LXB_4_ treatment inhibits transition to a functional reactive phenotype or promotes restoration of the microglia homeostatic phenotype.

**Figure 5:**
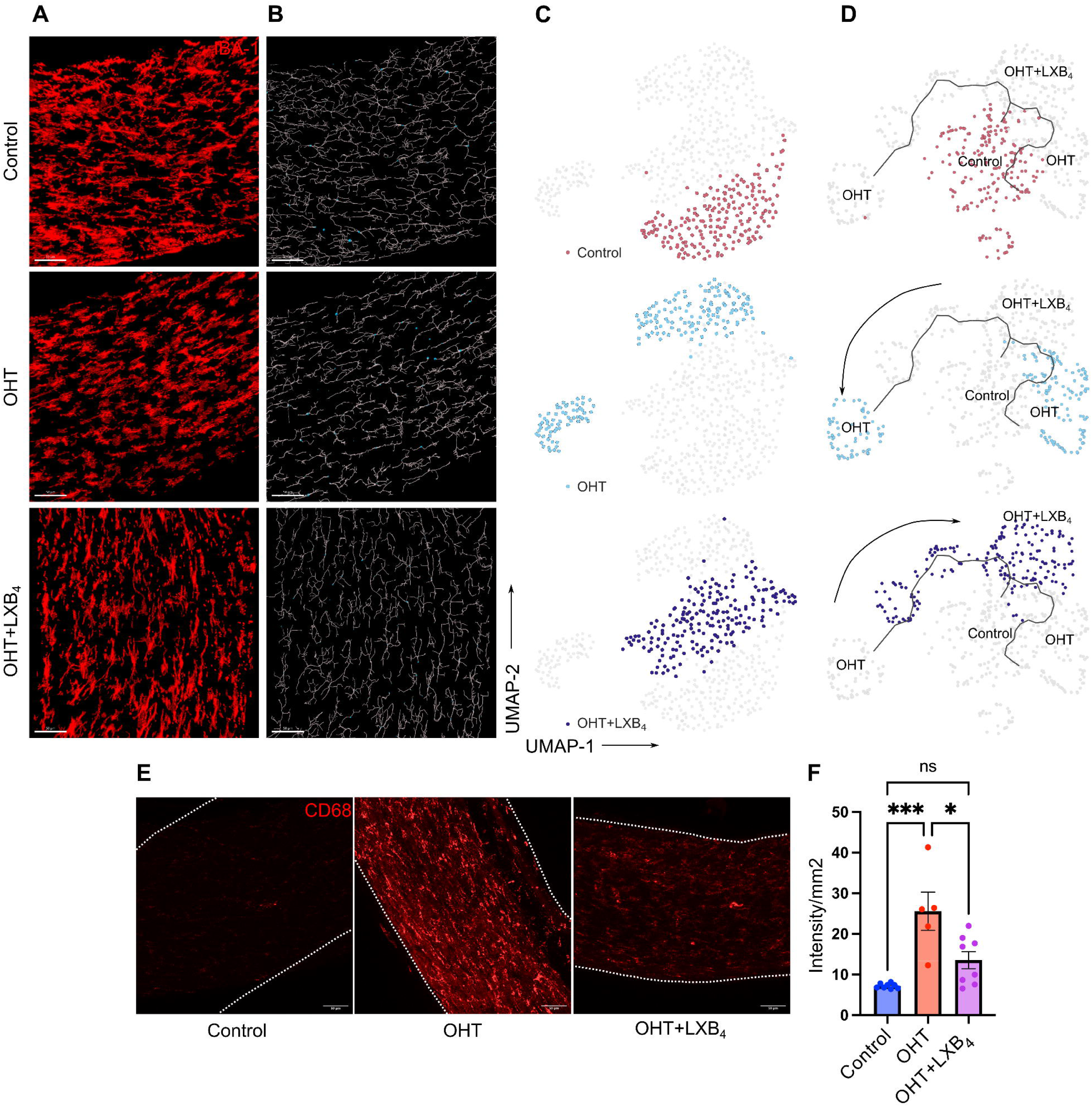
Neuroprotective LXB_4_ treatment induces a microglia homeostatic phenotype. **(A)** Representative confocal micrograph of Iba1 (red) stained optic nerve microglia in normotensive control, OHT, and OHT+LXB_4_ groups (scale bar-50μm). (**B)** Representative filament images of microglia morphology generated from the respective Iba1-stained images using IMARIS (scale bar-50μm). (**C)** UMAP plot displaying clustering of optic nerve microglia in normotensive control, OHT, and OHT+LXB_4_ groups. Each dot depicts 100 microglia bootstrapped for their topological morphology descriptor, and 200 dots are shown per group. (**D)** UMAP plot of pseudotime trajectory of microglia morphology for all groups. (**E)** Representative confocal micrographs of CD68-stained optic nerve microglia for normotensive control, OHT, and OHT+LXB_4_ (scale bar-50μm). (**F)** Quantification of CD68 expression in optic nerve whole-mounts for normotensive control (n=6), OHT (n=5), and OHT+LXB_4_ (n=8). Data were analyzed by One-way ANOVA with Tukey’s multiple comparison test (*p<0.05; ***p<0.001; ns, not significant). Each dot depicts the mean CD68 expression from a single retina. Data presented as mean ± SEM.

LXB_4_ intrinsic regulation of microglia homeostatic phenotype in the optic nerve was confirmed using Alox5 knockout (KO) mice, an established lipoxin-deficient mouse strain[50–52], in which the required enzyme (5-LOX) for lipoxin formation has been deleted. MorphOMICs analysis established that the microglia population in optic nerves from normotensive 5-LOX KO mice was distinct from that in healthy congenic wild-type mice (Supplementary Figure 7A). This suggests an altered homeostatic microglia phenotype in the absence of optic nerve homeostatic 5-LOX activity, i.e., lipoxin signaling. Consistent with the results from healthy optic nerves, morphOMICs and pseudotime trajectory analysis of optic nerve microglia after 1 wk of severe OHT showed that reactive microglia population in 5-LOX KO mice were distinct from wild-type congenic mice and farthest removed from the homeostatic microglia phenotype of congenic normotensive wild-type mice (Supplementary Figure 7B). These results suggest that the deletion of the lipoxin pathway in the optic nerve is associated with a dysregulated microglia phenotype and response to retinal OHT stress.

### LXB_4_ targets a marker of disease-associated microglia in the optic nerve

To investigate the mechanisms underlying LXB_4_ regulation of optic nerve microglia functional phenotypes during severe OHT, we conducted bulk RNA transcriptomics (RNA-seq) analysis of optic nerves from mice treated with or without LXB_4_. The myelinated optic nerve is comprised of a heterogeneous population of astrocytes, microglia, and oligodendrocytes. Hence, transcriptomic signatures from all these cell types[53] was expected. A linear regression model (DESeq2) was used to perform differential gene expression analysis comparing OHT versus normotensive controls (Figure 6A) and OHT with LXB_4_ treatment versus OHT (Figure 6B). Since some of microglia sensome genes were upregulated by moderate OHT (Figure 1F, G), we were interested in analyzing the expression of the large family of microglia sensome genes, which has been defined by Hickman et al. (2013)[38], in the optic nerve during severe OHT.

**Figure 6:**
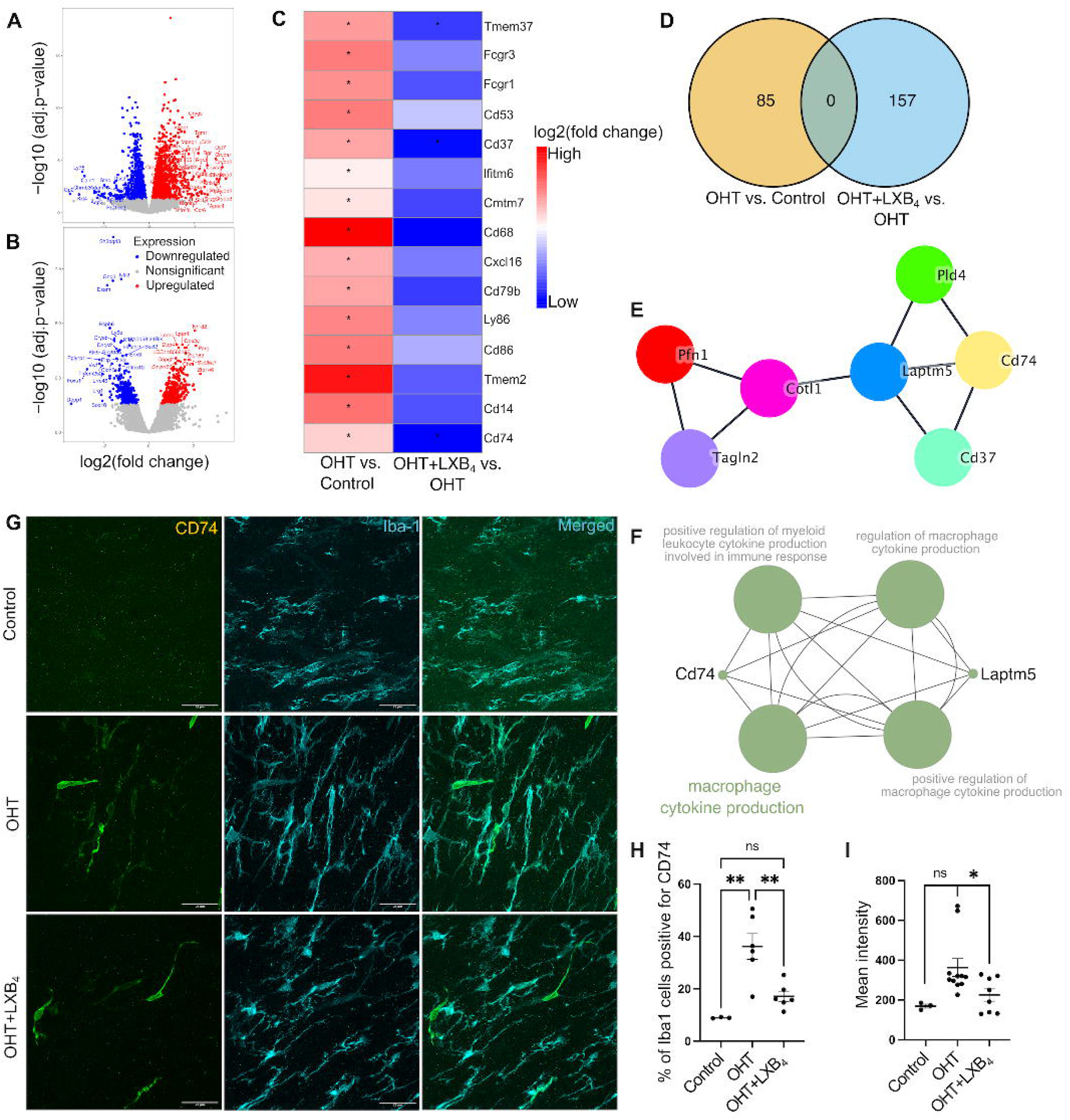
LXB_4_ regulates OHT induction of a CD74^+^ disease-associated microglia phenotype in the optic nerve. **(A)** Volcano plot for differentially expressed genes in the optic nerve between OHT versus normotensive control. (B) Volcano plot for differentially expressed genes in the optic nerve between OHT+LXB_4_ versus OHT (Genes: upregulated-red; downregulated-blue; no significant change-grey). (**C)** Heatmap showing expression of significantly changed (p.adjust<0.05) sensome genes in independent comparison of OHT versus normotensive control and OHT+LXB_4_ versus OHT; no Log_2_FC cutoff was applied. (**D)** Venn diagram for unique downregulated (Log_2_FC<-1, p.adjust<0.05) differentially expressed genes across independent comparisons of OHT versus normotensive control and OHT+LXB_4_ versus OHT. (**E)** Network of highly interacting genes in the optic nerve that were downregulated by LXB_4_ treatment during OHT, derived from MCODE analysis of STRING network. (**F)** Pathway enrichment network of genes shown in (E). **(G)** Representative confocal micrographs of optic nerve sections stained for Iba1 (cyan) and CD74 (red) in normotensive control, OHT, and OHT+LXB_4_ groups (scale bar-25 μm). (**H)** Quantification of Iba1 cells positive for CD74 expression (CD74^+^) in normotensive control (n=3), OHT (n=6), and OHT+LXB_4_ (n=6). Data were analyzed by One-way ANOVA with Tukey’s multiple comparisons test (**p<0.01; ns, not significant). Each dot depicts the mean measurement from a single optic nerve. (**I)** Quantification of CD74 expression intensity in normotensive control (n=3), OHT (n=11), and OHT+LXB_4_ (n=8). Data were analyzed by unpaired t-tests between normotensive control versus OHT and OHT versus OHT+LXB_4_ (*p<0.05; ns, not significant). Each dot depicts the mean measurement from a single optic nerve. Data presented as mean ± SEM.

Microglia sensome markers were significantly upregulated (*p.adjust<0.05) in the OHT versus control group (Figure 6C). Consistent with the homeostatic action of LXB_4_ in healthy retinas (Figure 1), LXB_4_ treatment during OHT significantly downregulated (*p.adjust<0.05) a subset of sensome markers in the optic nerve (Figure 6C). Given the role of LXB_4_ in regulating microglia phenotype, we focused our analysis on uniquely downregulated genes by LXB_4_ treatment during OHT (Figure 6D). STRING[54] protein-protein association clustering of genes downregulated by LXB_4_ treatment (Log2FC<-1, p.adjust<0.05) was followed by MCODE[55] analysis to identify highly interactive gene subclusters (Figure 6E). Pathway enrichment analysis of genes in these interactive subclusters revealed an association with *Cd74*, which regulates cytokine production in mononuclear phagocytes (Figure 6F). CD74 is a key marker for a unique subset of disease-associated microglia [56,57] that have been identified in central nervous system pathology[58]. *CD74* is detected at the RNA level in the retina and its RNA expression is increased in the optic nerve crush model[59]. To validate our findings, expression of CD74 was assessed by IHC in optic nerve sections from mice with OHT and mice with OHT that were treated with LXB_4_. CD74 expression was increased by 303.68% (p=0.0017) in Iba1-positive microglia cells in response to OHT (Figure 6G, H). LXB_4_ treatment markedly reduced the number of CD74-expressing cells by 52.56% (p=0.0054) (Figure 6G, H). Furthermore, the intensity of CD74 protein expression in the optic nerve was downregulated by 37.96% (p=0.033) following LXB_4_ treatment (Figure 6I). To assess the impact of endogenous LXB_4_ on CD74 in the optic nerve, protein expression was assessed by IHC in 5-LOX KO mice with and without severe OHT. Strikingly, optic nerves from normotensive 5-LOX KO mice showed a 468.9% (p= 0.0272) increase in the number Iba1 cells positive for CD74 compared to wild-type congenic control optic nerves (Supplementary Figure 8). OHT in 5-LOX KO mice did not further increase the number of Iba1 cells positive for CD74 in the optic nerves. However, the number of CD74 positive Iba1 cells in optic nerves was 503.3% (p= 0.0149) higher in 5-LOX KO mice when directly compared to wild-type congenic controls with OHT (Supplementary Figure 8). Surprisingly, CD74 expressing microglia were only detected by IHC in the optic nerve of wild-type mice but not in the retina during severe OHT (Supplementary Figure 9). These results provide evidence that OHT selectively induces a disease-associated microglia phenotype in the distal myelinated optic nerve and that LXB_4_ regulates these disease-associated microglia phenotype.

### LXB_4_ potentially regulates CD74^+^ disease-associated microglia and the phosphoinositide 3-kinase (PI3K) signaling pathway

Results revealed that LXB_4_ could regulate the functional phenotype of optic nerve microglia and downregulate the expression of CD74, a marker of disease-associated microglia in the brain. Subsequently, we investigated the intracellular signaling mechanisms underlying the regulation of disease-associated microglia in the optic nerve. KEGG pathway enrichment[60] analysis of genes downregulated (Log2FC <-1, p.adjust< 0.05) in the optic nerve by LXB_4_ treatment identified the PI3K-Akt signaling pathway as a significant candidate (Figure 7A). These results were of interest because PI3K signaling has been implicated in neuroinflammation and reactive glial phenotypes[61–63]. Considering that PI3K signaling is ubiquitously present in most cells[64], we performed co-staining of optic nerve sections with antibodies targeting CD74 and activated PI3K (phospho-PI3K, p-PI3K) to examine their expression specifically in the disease-associated microglia phenotype (Figure 7B). Initial analysis suggested co-expression of both p-PI3K and CD74 in many cells (Supplementary Figure 10A, B). Because of the membrane-binding nature of CD74 and the intracellular localization of p-PI3K, background noise, and expression of p-PI3K in other cells, it was challenging to perform co-localization and expression analysis comparisons between control and severe OHT plus or minus LXB_4_ treatment.

**Figure 7:**
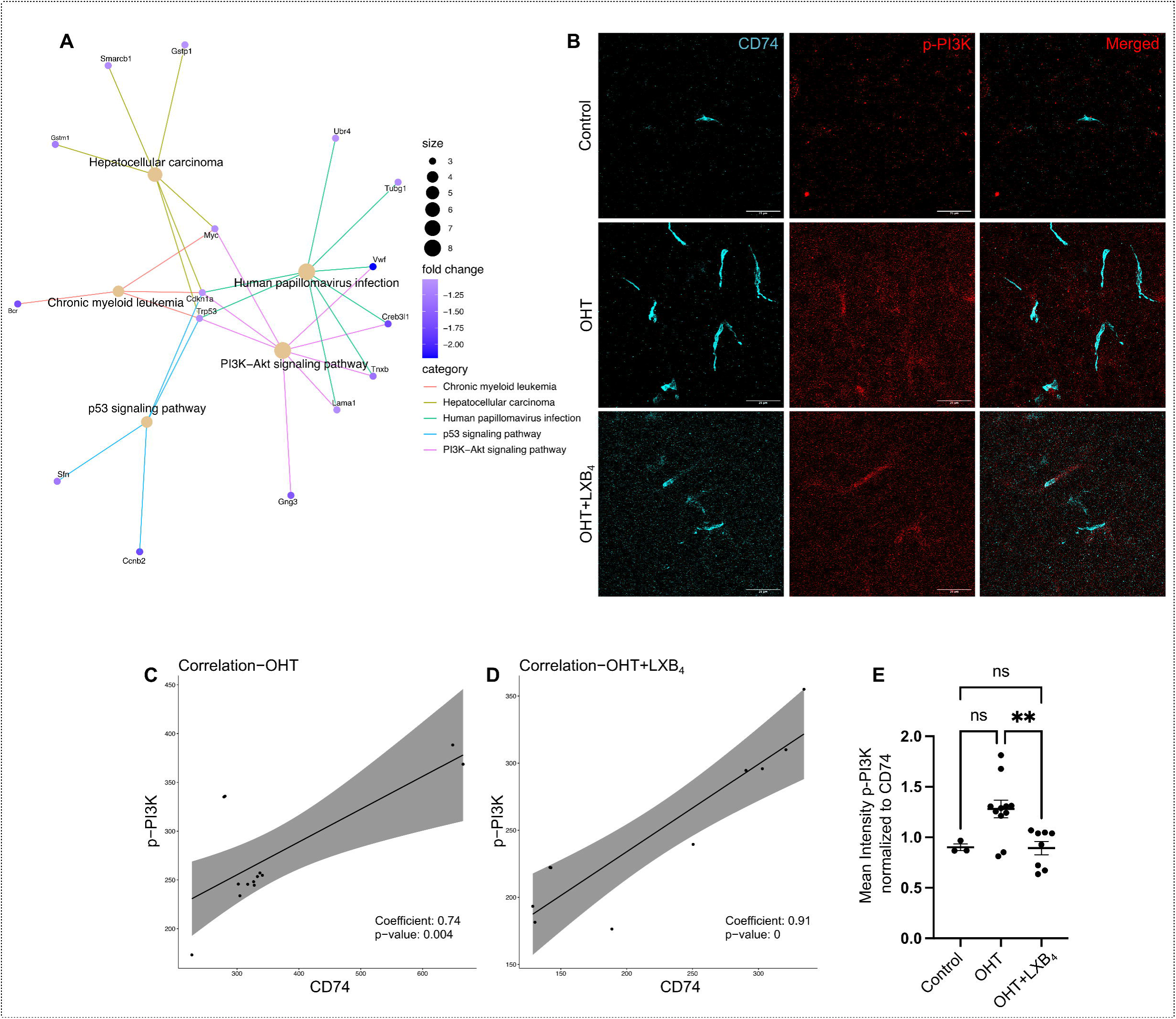
LXB_4_ regulates PI3K signaling in optic nerve CD74^+^ disease-associated microglia. **(A)** KEGG pathway enrichment map of genes downregulated by LXB_4_ treatment during severe OHT. The Pathway circle size depicts number of genes involved in enrichment; genes linked to pathways are shown with their fold change scale. (**B)** Representative confocal micrographs of optic nerve sections stained for CD74 (cyan) and p-PI3K (red) in normotensive control, OHT, and OHT+LXB_4_ groups (scale bar-25 μm). (**C, D)** Correlation plot for CD74 and p-PI3K protein expression in OHT (**C)** and OHT+LXB_4_ (**D).** Each dot represents the intensity of CD74 and p-PI3K correlated from a single cell. (**E)** Quantification of normalized p-PI3K to CD74 expression. Data were analyzed One-way ANOVA with Tukey’s multiple comparison test (**p<0.01; ns, not significant). Each dot represents normalized expression intensity from a single cell. Data presented as mean ± SEM.

Therefore, to establish the regulatory role of PI3K signaling in disease-associated microglia, correlation analysis was used to examine the expression of these markers in the optic nerve. Correlation analysis is a well-established approach for determining regulatory relationships[65]. Significant positive correlation was identified between p-PI3K and CD74 expression in both the OHT (r= 0.74, p=0.004, 95% confidence interval= [0.31,0.92]) (Figure 7C) and OHT with LXB_4_ treatment groups (r=0.91, p=0, 95% confidence interval= [0.65,0.98]) (Figure 7D). Correlation analysis of CD74 and p-PI3K with DAPI signals (nuclear staining in all cells) was used as the negative correlation control. As expected, DAPI staining did not correlate with p-PI3K (r= 0.34, p=0.258, 95% confidence interval= [−0.26,0.75]) (Supplementary Figure 10C) or CD74 (r= 0.51, p=0.077, 95% confidence interval= [−0.06,0.83]) (Supplementary Figure 10D).

Given the consistent correlation between CD74 and p-PI3K co-expression in the optic nerves of mice with OHT, it is of interest that LXB_4_ treatment significantly downregulated p-PI3K relative to CD74 (30.4%, p=0.0072) (Figure 7E). Taken together, the correlation analysis suggests that activated PI3K during OHT is a feature of the optic nerve disease-associated (CD74) microglia phenotype. The mechanism for LXB_4_ downregulation of a CD74 microglia phenotype during OHT could potentially be mediated by inhibition of PI3K signaling.

## Discussion

The current study identifies microglia as a target for regulation by LXB_4_ in the retina and myelinated, retrolaminar optic nerve. Microglia are specialized and tissue resident mononuclear phagocytes that are essential for maintaining retinal and brain homeostasis but are also a key cell type for driving pathogenesis in neurodegenerative diseases such as Alzheimer’s and glaucoma[22–24,66]. Our previous study[6] established endogenous LXB_4_ formation in the retina and optic nerve head. More importantly, it demonstrated direct regulation of RGCs as a cellular target in the retina for LXB_4_’s neuroprotective action in both retinal excitotoxicity and OHT injury. The unexpected finding that regulation of the microglia pathway for environmental sensing and transition to a reactive phenotype are primary targets for LXB_4_ in the healthy retina is of great interest. Changes in the microglia sensome are key events in neurodegenerative diseases and drive the transition of the microglia phenotype from a neuroprotective to a neurotoxic state[38]. Neurotoxic microglia activate astrocytic reactivity in the brain[23] and sustained activation of microglia is related to the degeneration of RGCs in glaucoma models[67,68]. Hence, the regulation of microglial sensome by LXB_4_ is likely a mechanism for its homeostatic and neuroprotective role in the retina and optic nerve.

Knowledge of the mechanisms of action and cellular targets of LXB_4_ is limited, despite several publications establishing endogenous formation of LXB_4_ and its ability to counter-regulate the activity of pro-inflammatory signals in leukocytes during acute inflammation[9,69]. An early study[70] identified a unique ability of LXB_4_ to drive migration and adhesion of monocytes, the mononuclear phagocytes in blood, without activating an inflammatory monocyte phenotype. Our *in vivo* findings indicate that LXB_4_ regulation of pathways essential to controlling transition to a reactive inflammatory phenotype as a potential mechanism for maintaining the homeostatic functions of the resident mononuclear phagocyte in central nervous system, namely microglia. Interestingly, the structurally and functionally distinct LXA_4_, which acts at a different receptor than LXB_4,_ has been found to inhibit the inflammatory microglial function in spinal cord injury[71] and in severe congenital retinal degeneration[72] mouse models. However, it is important to point out that even though both LXA_4_ and LXB_4_ are generated in the healthy retina and optic nerve, they act on different receptors and LXB_4_ is the most potent lipoxin in terms of *in vivo* and *in vitro* neuroprotective activity[6,11]. We previously reported that glaucomatous injury downregulates enzymes required for LXB_4_ formation in the retina[6], which underscores the homeostatic role of lipoxin in the retina. This finding is consistent with a subsequent report in the RD1 mouse strain, which develops severe photoreceptor death and retinal degeneration that correlates with marked downregulation of the lipoxin pathway[72].

An important finding of the study is that both severe and moderate OHT trigger dynamic and temporally defined morphological changes in the distal optic nerve microglia population that paralleled the temporal morphological response of the retinal microglia population. Microglia populations in both the retina and optic nerve transitioned to morphologically distinct populations during the time course of sustained long-term OHT. Interestingly, after polarizing towards a functional phenotype that progressively is more divergent from the homeostatic population in normotensive condition, microglia transition back towards a homeostatic morphology by week 6, despite unchanged and sustained OHT, suggesting an adaptive response. The behavioral response of optic nerve microglia provided compelling evidence that the myelinated optic nerve rapidly responds to OHT in the retina. In addition, the possibility of macrophage infiltration in the optic nerve cannot be excluded since CD74 and/or Iba1 can also be expressed in macrophages. However, the total number of Iba1 positive cells in the optic nerve did not change during OHT suggesting that there was no proliferation of microglia or infiltration of macrophages. Research to understand the pathogenesis of glaucomatous RGC degeneration has primarily focused on the retina and optic nerve head. However, emerging evidence points toward RGC axons as a primary site of glaucoma pathogenesis, including impaired axonal transport and mitochondrial dysfunction as important early events that precede the loss of RGC in the retina[73,74]. At this time, it is unclear how microglia in the distal myelinated optic nerve sense and rapidly respond to OHT. It is tempting to speculate that abnormal axonal transport and energy metabolism are sensed by the abundant resident optic nerve microglia population.

A key finding is that OHT leads to expression of a unique CD74 positive microglia population in the optic nerve. This population was unique to the optic nerve, and CD74 positive microglia were not present in the retina during moderate or severe OHT. CD74 positive microglia have been defined as a disease-associated microglia population (DAM) in the brain. DAM were initially identified in a mouse model of Alzheimer’s disease (AD) using single-cell transcriptomics and proteomics analysis[57,75–81]. Subsequently, their association with the disease was confirmed through their presence in affected brain regions and human genome-wide association studies[56]. The regulation of DAM, particularly under neuropathic conditions is of great interest and a potential therapeutic target. DAM activation requires two stage TREM2 mediated signaling which results in upregulation of phagocytic, lysosomal and lipid metabolism pathways[56]. Understanding the role of DAM in neuropathic conditions is challenging because TREM2 plays distinct functional roles at different stages of AD pathology[82].

Hence, it is of great interest that neuroprotective LXB_4_ treatment inhibited the induction of an optic nerve CD74 microglia population in response to OHT, in addition to inhibiting microglia reactivity (CD68 expression). LXB_4_ treatment correlated with a distinct functional microglia phenotype compared to untreated mice with OHT. More importantly, LXB_4_ treatment resulted in a microglial phenotype that was morphologically closer to the healthy homeostatic phenotype compared to untreated mice. LXB_4_ has established direct action with other mononuclear phagocytes[70], namely monocytes and macrophages, but it is unclear if the regulation of microglia functional responses in the optic nerve by LXB_4_ treatment is due to direct regulation or potentially through indirect pathways involving other retinal cells[6,37]. In addition, the relevance of the endogenous lipoxin pathway in regulating microglia homeostatic function is underscored by the loss of function in 5-LOX KO mice. This lipoxin-deficient mouse line exhibited presence of CD74 positive microglia in the optic nerve without OHT, which was equivalent to congenic wild-type mice with OHT. In addition, morphOMIC analysis established that the homeostatic optic nerve microglia population was distinct in 5-LOX KO mice compared to their healthy congenic wild-type controls and exhibited an amplified reactive response to OHT. 5-LOX is also a key enzyme for generating pro-inflammatory leukotrienes; hence, the phenotype of an amplified optic nerve DAM and reactive microglia response is unexpected. However, we have not detected significant leukotriene generation in mouse or rat healthy or injured retinas or optic nerves[6,51,83]. More importantly, 5-LOX inhibition or deletion amplifies retinal inflammation[83], autoimmune[51] and neurodegeneration pathogenesis[6] indicating that 5-LOX has protective functions in the retina.

Pathway analysis and IHC established that OHT correlated with an increase in PI3K activity in optic nerve microglia, including the unique CD74 positive microglia population. More importantly, downregulation of the CD74 microglia population by LXB_4_ treatment correlated with reduced PI3 kinase activity and expression in the optic nerve. These findings suggest that inhibition of the PI3K pathway is a potential target for LXB_4_ signaling. The PI3K pathway is expressed in many cell types with broad cell-specific functions[84]. Microglia activation of the PI3K-Akt pathway increases the formation and secretion of pro-inflammatory cytokines such as IL-6, IL-1β, IL-12, and TNF-α, which drive neurodegeneration[61–63], activate NF-kB, and induce pro-inflammatory mediators in *in vitro* microglia models of LPS-induced inflammation[61,85]. Inhibition of PI3K/Akt signaling has been shown to protect RGCs after optic nerve injury in rat models[86]. PI3K signaling also provides pro-proliferative and pro-survival functions towards sustained microglia activation via TREM2 regulation[56,87], and hence becomes an important mediator in regulating DAM. The canonical anti-inflammatory action of LXB_4_ is the inhibition of cytokine production by leukocytes[9,88], which is consistent with our finding that LXB_4_ treatment inhibits PI3K activation in a reactive microglial phenotype.

In conclusion, our study reveals that OHT triggers early and dynamic functional microglial responses and the presence of a unique disease-associated microglia phenotype in the distal myelinated optic nerve. Treatment with neuroprotective LXB_4_ regulates microglia homeostatic function by inhibiting transition to a functional reactive phenotype or promoting restoration of the microglia homeostatic phenotype. Of particular interest is the finding that LXB_4_ treatment and the endogenous LXB_4_ pathway inhibit microglia polarization to a CD74 disease-associated phenotype in the optic nerve. These findings indicate that the regulation of microglia functions as a potential mechanism for LXB_4_-mediated neuroprotection and dynamic optic nerve microglia responses which is an unexpected early event in retinal pathogenesis following OHT.

## Declarations

### Ethics approval and consent to participate

All animals were handled in accordance with the ARVO Statement for the Use of Animals in Ophthalmic and Vision Research, and all procedures were approved by the Institutional Animal Care and Use Committee (IACUC) at the University of California, Berkeley.

### Consent for publication

Not applicable

### Availability of data and materials

The dataset generated in this study was deposited to the GEO (GSE251716, scRNA-seq; GSE250615, bulk RNA-seq), and is publicly available at the publication date of this article.

### Authors’ contributions

S.M., J.S., J.G.F, and K.G. conceived the study; S.M., M.L., S.K., T.S., and M.K. performed the experiments; S.M. and M.L. performed data analysis; S.M., E.W., J.G.F., and K.G. wrote the manuscript.

## Acknowledgments

We acknowledge the QB3 Genomics core RRID: SCR_022170, UC Berkeley, for sequencing support, and The CNR Biological Imaging Facility, CRL Molecular Imaging Center, RRID: SCR_017852, UC Berkeley, for imaging and image analysis support.

## References

1. Calkins DJ, Pekny M, Cooper ML, Benowitz L, Lasker/IRRF Initiative on Astrocytes and Glaucomatous Neurodegeneration Participants. The challenge of regenerative therapies for the optic nerve in glaucoma. Exp Eye Res. 2017;157:28–33.

2. Alqawlaq S, Flanagan JG, Sivak JM. All roads lead to glaucoma: Induced retinal injury cascades contribute to a common neurodegenerative outcome. Exp Eye Res. 2019;183:88–97.

3. Weinreb RN, Aung T, Medeiros FA. The Pathophysiology and Treatment of Glaucoma. JAMA. 2014;311:1901–11.

4. Wareham LK, Liddelow SA, Temple S, Benowitz LI, Di Polo A, Wellington C, et al. Solving neurodegeneration: common mechanisms and strategies for new treatments. Molecular Neurodegeneration. 2022;17:23.

5. Almasieh M, Wilson AM, Morquette B, Cueva Vargas JL, Di Polo A. The molecular basis of retinal ganglion cell death in glaucoma. Prog Retin Eye Res. 2012;31:152–81.

6. Livne-Bar I, Wei J, Liu H-H, Alqawlaq S, Won G-J, Tuccitto A, et al. Astrocyte-derived lipoxins A4 and B4 promote neuroprotection from acute and chronic injury. J Clin Invest. 2017;127:4403–14.

7. Wei J, Gronert K. Eicosanoid and Specialized Proresolving Mediator Regulation of Lymphoid Cells. Trends Biochem Sci. 2019;44:214–25.

8. Wei J, Gronert K. The Role of Pro-resolving Lipid Mediators in Ocular Diseases. Mol Aspects Med. 2017;58:37–43.

9. Serhan CN. Pro-resolving lipid mediators are leads for resolution physiology. Nature. 2014;510:92–101.

10. Flitter BA, Fang X, Matthay MA, Gronert K. The potential of lipid mediator networks as ocular surface therapeutics and biomarkers. Ocul Surf. 2021;19:104–14.

11. Kim C, Livne-Bar I, Gronert K, Sivak JM. Fairweather Friends: Evidence of Lipoxin Dysregulation in Neurodegeneration. Mol Nutr Food Res. 2020;64:e1801076.

12. Romano M, Maddox JF, Serhan CN. Activation of human monocytes and the acute monocytic leukemia cell line (THP-1) by lipoxins involves unique signaling pathways for lipoxin A4 versus lipoxin B4: evidence for differential Ca2+ mobilization. J Immunol. 1996;157:2149–54.

13. Heuss ND, Pierson MJ, Roehrich H, McPherson SW, Gram AL, Li L, et al. Optic nerve as a source of activated retinal microglia post-injury. Acta Neuropathologica Communications. 2018;6:66.

14. Neufeld AH. Microglia in the optic nerve head and the region of parapapillary chorioretinal atrophy in glaucoma. Arch Ophthalmol. 1999;117:1050–6.

15. Ahmad I, Subramani M. Microglia: Friends or Foes in Glaucoma? A Developmental Perspective. Stem Cells Translational Medicine. 2022;11:1210–8.

16. Au NPB, Ma CHE. Neuroinflammation, Microglia and Implications for Retinal Ganglion Cell Survival and Axon Regeneration in Traumatic Optic Neuropathy. Front Immunol. 2022;13:860070.

17. Guo L, Choi S, Bikkannavar P, Cordeiro MF. Microglia: Key Players in Retinal Ageing and Neurodegeneration. Front Cell Neurosci. 2022;16:804782.

18. Hickman S, Izzy S, Sen P, Morsett L, El Khoury J. Microglia in neurodegeneration. Nat Neurosci. 2018;21:1359–69.

19. Zhao X, Sun R, Luo X, Wang F, Sun X. The Interaction Between Microglia and Macroglia in Glaucoma. Frontiers in Neuroscience. 2021;15:573.

20. Fernández-Albarral JA, Ramírez AI, de Hoz R, Salazar JJ. Retinal microglial activation in glaucoma: evolution over time in a unilateral ocular hypertension model. Neural Regen Res. 2021;17:797–9.

21. Zeng H-L, Shi J-M. The role of microglia in the progression of glaucomatous neurodegeneration-a review. Int J Ophthalmol. 2018;11:143–9.

22. Jha MK, Jo M, Kim J-H, Suk K. Microglia-Astrocyte Crosstalk: An Intimate Molecular Conversation. Neuroscientist. 2019;25:227–40.

23. Liddelow SA, Guttenplan KA, Clarke LE, Bennett FC, Bohlen CJ, Schirmer L, et al. Neurotoxic reactive astrocytes are induced by activated microglia. Nature. 2017;541:481–7.

24. Joshi AU, Minhas PS, Liddelow SA, Haileselassie B, Andreasson KI, Dorn GW, et al. Fragmented mitochondria released from microglia trigger A1 astrocytic response and propagate inflammatory neurodegeneration. Nat Neurosci. 2019;22:1635–48.

25. Zhang J, Li L, Huang H, Fang F, Webber HC, Zhuang P, et al. Silicone oil-induced ocular hypertension and glaucomatous neurodegeneration in mouse. Nathans J, Calabrese RL, Ou Y, editors. eLife. 2019;8:e45881.

26. Fang F, Zhang J, Zhuang P, Liu P, Li L, Huang H, et al. Chronic mild and acute severe glaucomatous neurodegeneration derived from silicone oil-induced ocular hypertension. Sci Rep. 2021;11:9052.

27. Liu H-H, Zhang L, Shi M, Chen L, Flanagan JG. Comparison of laser and circumlimbal suture induced elevation of intraocular pressure in albino CD-1 mice. PLoS One. 2017;12:e0189094.

28. Liu H-H, Flanagan JG. A Mouse Model of Chronic Ocular Hypertension Induced by Circumlimbal Suture. Invest Ophthalmol Vis Sci. 2017;58:353–61.

29. Heindl S, Gesierich B, Benakis C, Llovera G, Duering M, Liesz A. Automated Morphological Analysis of Microglia After Stroke. Front Cell Neurosci. 2018;12:106.

30. Qiu X, Mao Q, Tang Y, Wang L, Chawla R, Pliner HA, et al. Reversed graph embedding resolves complex single-cell trajectories. Nat Methods. 2017;14:979–82.

31. Cao J, Spielmann M, Qiu X, Huang X, Ibrahim DM, Hill AJ, et al. The single-cell transcriptional landscape of mammalian organogenesis. Nature. 2019;566:496–502.

32. Trapnell C, Cacchiarelli D, Grimsby J, Pokharel P, Li S, Morse M, et al. The dynamics and regulators of cell fate decisions are revealed by pseudotemporal ordering of single cells. Nat Biotechnol. 2014;32:381–6.

33. Hao Y, Hao S, Andersen-Nissen E, Mauck WM, Zheng S, Butler A, et al. Integrated analysis of multimodal single-cell data. Cell. 2021;184:3573–3587.e29.

34. Tran NM, Shekhar K, Whitney IE, Jacobi A, Benhar I, Hong G, et al. Single-Cell Profiles of Retinal Ganglion Cells Differing in Resilience to Injury Reveal Neuroprotective Genes. Neuron. 2019;104:1039–1055.e12.

35. Peng Y-R, Shekhar K, Yan W, Herrmann D, Sappington A, Bryman GS, et al. Molecular Classification and Comparative Taxonomics of Foveal and Peripheral Cells in Primate Retina. Cell. 2019;176:1222–1237.e22.

36. Yan W, Laboulaye MA, Tran NM, Whitney IE, Benhar I, Sanes JR. Mouse Retinal Cell Atlas: Molecular Identification of over Sixty Amacrine Cell Types. J Neurosci. 2020;40:5177–95.

37. Karnam S, Maurya S, Ng E, Choudhary A, Thobani A, Flanagan JG, et al. Dysregulation of neuroprotective lipoxin pathway in astrocytes in response to cytokines and ocular hypertension. Acta Neuropathol Commun. 2024;12:58.

38. Hickman SE, Kingery ND, Ohsumi T, Borowsky M, Wang L, Means TK, et al. The Microglial Sensome Revealed by Direct RNA Sequencing. Nat Neurosci. 2013;16:1896–905.

39. Tan Z, Guo Y, Shrestha M, Sun D, Gregory-Ksander M, Jakobs TC. Microglia depletion exacerbates retinal ganglion cell loss in a mouse model of glaucoma. Exp Eye Res. 2022;225:109273.

40. Zhang J, Fang F, Li L, Huang H, Webber HC, Sun Y, et al. A Reversible Silicon Oil-Induced Ocular Hypertension Model in Mice. J Vis Exp. 2019;

41. Zhang J, Hu Y. Comparing silicone oil-induced ocular hypertension with other inducible glaucoma models in mice. Neural Regen Res. 2020;15:1652–3.

42. Sousa C, Golebiewska A, Poovathingal SK, Kaoma T, Pires-Afonso Y, Martina S, et al. Single-cell transcriptomics reveals distinct inflammation-induced microglia signatures. EMBO reports. 2018;19:e46171.

43. Guergues J, Wohlfahrt J, Zhang P, Liu B, Stevens Jr. SM. Deep proteome profiling reveals novel pathways associated with pro-inflammatory and alcohol-induced microglial activation phenotypes. Journal of Proteomics. 2020;220:103753.

44. Wang J, He W, Zhang J. A richer and more diverse future for microglia phenotypes. Heliyon. 2023;9:e14713.

45. Schwabenland M, Brück W, Priller J, Stadelmann C, Lassmann H, Prinz M. Analyzing microglial phenotypes across neuropathologies: a practical guide. Acta Neuropathol. 2021;142:923–36.

46. Lier J, Streit WJ, Bechmann I. Beyond Activation: Characterizing Microglial Functional Phenotypes. Cells. 2021;10:2236.

47. Colombo G, Cubero RJA, Kanari L, Venturino A, Schulz R, Scolamiero M, et al. A tool for mapping microglial morphology, morphOMICs, reveals brain-region and sex-dependent phenotypes. Nat Neurosci. 2022;25:1379–93.

48. Kanari L, Dłotko P, Scolamiero M, Levi R, Shillcock J, Hess K, et al. A Topological Representation of Branching Neuronal Morphologies. Neuroinform. 2018;16:3–13.

49. Simpson JE, Ince PG, Higham CE, Gelsthorpe CH, Fernando MS, Matthews F, et al. Microglial activation in white matter lesions and nonlesional white matter of ageing brains. Neuropathol Appl Neurobiol. 2007;33:670–83.

50. Leedom AJ, Sullivan AB, Dong B, Lau D, Gronert K. Endogenous LXA4 circuits are determinants of pathological angiogenesis in response to chronic injury. Am J Pathol. 2010;176:74–84.

51. Wei J, Mattapallil MJ, Horai R, Jittayasothorn Y, Modi AP, Sen HN, et al. A novel role for lipoxin A4 in driving a lymph node–eye axis that controls autoimmunity to the neuroretina. Taniguchi T, Smith L, Bazan NG, editors. eLife. 2020;9:e51102.

52. Prieto P, Cuenca J, Través PG, Fernández-Velasco M, Martín-Sanz P, Boscá L. Lipoxin A4 impairment of apoptotic signaling in macrophages: implication of the PI3K/Akt and the ERK/Nrf-2 defense pathways. Cell Death Differ. 2010;17:1179–88.

53. Pan Y-B, Sun Y, Li H-J, Zhou L-Y, Zhang J, Feng D-F. Transcriptome Analyses Reveal Systematic Molecular Pathology After Optic Nerve Crush. Front Cell Neurosci. 2022;15:800154.

54. Szklarczyk D, Kirsch R, Koutrouli M, Nastou K, Mehryary F, Hachilif R, et al. The STRING database in 2023: protein-protein association networks and functional enrichment analyses for any sequenced genome of interest. Nucleic Acids Res. 2023;51:D638–46.

55. Bader GD, Hogue CWV. An automated method for finding molecular complexes in large protein interaction networks. BMC Bioinformatics. 2003;4:2.

56. Deczkowska A, Keren-Shaul H, Weiner A, Colonna M, Schwartz M, Amit I. Disease-Associated Microglia: A Universal Immune Sensor of Neurodegeneration. Cell. 2018;173:1073–81.

57. Keren-Shaul H, Spinrad A, Weiner A, Matcovitch-Natan O, Dvir-Szternfeld R, Ulland TK, et al. A Unique Microglia Type Associated with Restricting Development of Alzheimer’s Disease. Cell. 2017;169:1276–1290.e17.

58. Olah M, Menon V, Habib N, Taga MF, Ma Y, Yung CJ, et al. Single cell RNA sequencing of human microglia uncovers a subset associated with Alzheimer’s disease. Nat Commun. 2020;11:6129.

59. Templeton JP, Freeman NE, Nickerson JM, Jablonski MM, Rex TS, Williams RW, et al. Innate immune network in the retina activated by optic nerve crush. Invest Ophthalmol Vis Sci. 2013;54:2599–606.

60. Kanehisa M, Furumichi M, Tanabe M, Sato Y, Morishima K. KEGG: new perspectives on genomes, pathways, diseases and drugs. Nucleic Acids Res. 2017;45:D353–61.

61. Saponaro C, Cianciulli A, Calvello R, Dragone T, Iacobazzi F, Panaro MA. The PI3K/Akt pathway is required for LPS activation of microglial cells. Immunopharmacol Immunotoxicol. 2012;34:858–65.

62. Cianciulli A, Porro C, Calvello R, Trotta T, Lofrumento DD, Panaro MA. Microglia Mediated Neuroinflammation: Focus on PI3K Modulation. Biomolecules. 2020;10:137.

63. London A, Cohen M, Schwartz M. Microglia and monocyte-derived macrophages: functionally distinct populations that act in concert in CNS plasticity and repair. Front Cell Neurosci. 2013;7:34.

64. Liu P, Cheng H, Roberts TM, Zhao JJ. Targeting the phosphoinositide 3-kinase (PI3K) pathway in cancer. Nat Rev Drug Discov. 2009;8:627–44.

65. Wu Y, Eghbali M, Ou J, Lu R, Toro L, Stefani E. Quantitative Determination of Spatial Protein-Protein Correlations in Fluorescence Confocal Microscopy. Biophys J. 2010;98:493–504.

66. Wei X, Cho K-S, Thee EF, Jager MJ, Chen DF. Neuroinflammation and microglia in glaucoma: time for a paradigm shift. J Neurosci Res. 2019;97:70–6.

67. Bosco A, Inman DM, Steele MR, Wu G, Soto I, Marsh-Armstrong N, et al. Reduced retina microglial activation and improved optic nerve integrity with minocycline treatment in the DBA/2J mouse model of glaucoma. Invest Ophthalmol Vis Sci. 2008;49:1437–46.

68. Fischer AJ, Zelinka C, Milani-Nejad N. Reactive retinal microglia, neuronal survival, and the formation of retinal folds and detachments. Glia. 2015;63:313–27.

69. Jaén RI, Sánchez-García S, Fernández-Velasco M, Boscá L, Prieto P. Resolution-Based Therapies: The Potential of Lipoxins to Treat Human Diseases. Front Immunol. 2021;12:658840.

70. Maddox JF, Serhan CN. Lipoxin A4 and B4 are potent stimuli for human monocyte migration and adhesion: selective inactivation by dehydrogenation and reduction. J Exp Med. 1996;183:137–46.

71. Martini AC, Berta T, Forner S, Chen G, Bento AF, Ji R-R, et al. Lipoxin A4 inhibits microglial activation and reduces neuroinflammation and neuropathic pain after spinal cord hemisection. J Neuroinflammation. 2016;13:75.

72. Lu Z, Zhang H, Zhang X, Gao Y, Yin ZQ. Lipoxin A4 delays the progression of retinal degeneration via the inhibition of microglial overactivation. Biochem Biophys Res Commun. 2019;516:900–6.

73. Jassim AH, Inman DM, Mitchell CH. Crosstalk Between Dysfunctional Mitochondria and Inflammation in Glaucomatous Neurodegeneration. Front Pharmacol. 2021;12:699623.

74. Quintero H, Shiga Y, Belforte N, Alarcon-Martinez L, El Hajji S, Villafranca-Baughman D, et al. Restoration of mitochondria axonal transport by adaptor Disc1 supplementation prevents neurodegeneration and rescues visual function. Cell Rep. 2022;40:111324.

75. Ajami B, Samusik N, Wieghofer P, Ho PP, Crotti A, Bjornson Z, et al. Single-cell mass cytometry reveals distinct populations of brain myeloid cells in mouse neuroinflammation and neurodegeneration models. Nat Neurosci. 2018;21:541–51.

76. Friedman BA, Srinivasan K, Ayalon G, Meilandt WJ, Lin H, Huntley MA, et al. Diverse Brain Myeloid Expression Profiles Reveal Distinct Microglial Activation States and Aspects of Alzheimer’s Disease Not Evident in Mouse Models. Cell Reports. 2018;22:832–47.

77. Holtman IR, Raj DD, Miller JA, Schaafsma W, Yin Z, Brouwer N, et al. Induction of a common microglia gene expression signature by aging and neurodegenerative conditions: a co-expression meta-analysis. acta neuropathol commun. 2015;3:31.

78. Kamphuis W, Kooijman L, Schetters S, Orre M, Hol EM. Transcriptional profiling of CD11c-positive microglia accumulating around amyloid plaques in a mouse model for Alzheimer’s disease. Biochimica et Biophysica Acta (BBA) - Molecular Basis of Disease. 2016;1862:1847–60.

79. Krasemann S, Madore C, Cialic R, Baufeld C, Calcagno N, El Fatimy R, et al. The TREM2-APOE Pathway Drives the Transcriptional Phenotype of Dysfunctional Microglia in Neurodegenerative Diseases. Immunity. 2017;47:566–581.e9.

80. Mrdjen D, Pavlovic A, Hartmann FJ, Schreiner B, Utz SG, Leung BP, et al. High-Dimensional Single-Cell Mapping of Central Nervous System Immune Cells Reveals Distinct Myeloid Subsets in Health, Aging, and Disease. Immunity. 2018;48:380–395.e6.

81. Ofengeim D, Mazzitelli S, Ito Y, DeWitt JP, Mifflin L, Zou C, et al. RIPK1 mediates a disease-associated microglial response in Alzheimer’s disease. Proceedings of the National Academy of Sciences. 2017;114:E8788–97.

82. Jay TR, Hirsch AM, Broihier ML, Miller CM, Neilson LE, Ransohoff RM, et al. Disease Progression-Dependent Effects of TREM2 Deficiency in a Mouse Model of Alzheimer’s Disease. J Neurosci. 2017;37:637–47.

83. Sapieha P, Stahl A, Chen J, Seaward MR, Willett KL, Krah NM, et al. 5-Lipoxygenase metabolite 4-HDHA is a mediator of the antiangiogenic effect of ω-3 polyunsaturated fatty acids. Sci Transl Med. 2011;3:69ra12.

84. He X, Li Y, Deng B, Lin A, Zhang G, Ma M, et al. The PI3K/AKT signalling pathway in inflammation, cell death and glial scar formation after traumatic spinal cord injury: Mechanisms and therapeutic opportunities. Cell Prolif. 2022;55:e13275.

85. Zhu Y, Chen X, Liu Z, Peng Y-P, Qiu Y-H. Interleukin-10 Protection against Lipopolysaccharide-Induced Neuro-Inflammation and Neurotoxicity in Ventral Mesencephalic Cultures. Int J Mol Sci. 2015;17:25.

86. Luo J-M, Cen L-P, Zhang X-M, Chiang SW-Y, Huang Y, Lin D, et al. PI3K/akt, JAK/STAT and MEK/ERK pathway inhibition protects retinal ganglion cells via different mechanisms after optic nerve injury. Eur J Neurosci. 2007;26:828–42.

87. Peng Q, Malhotra S, Torchia JA, Kerr WG, Coggeshall KM, Humphrey MB. TREM2- and DAP12-dependent activation of PI3K requires DAP10 and is inhibited by SHIP1. Sci Signal. 2010;3:ra38.

88. Chandrasekharan JA, Sharma-Walia N. Lipoxins: nature’s way to resolve inflammation. J Inflamm Res. 2015;8:181–92.

