## Supplementary File for "Regulation of Diseases-Associated Microglia in the Optic Nerve by Lipoxin B_4_ and Ocular Hypertension"

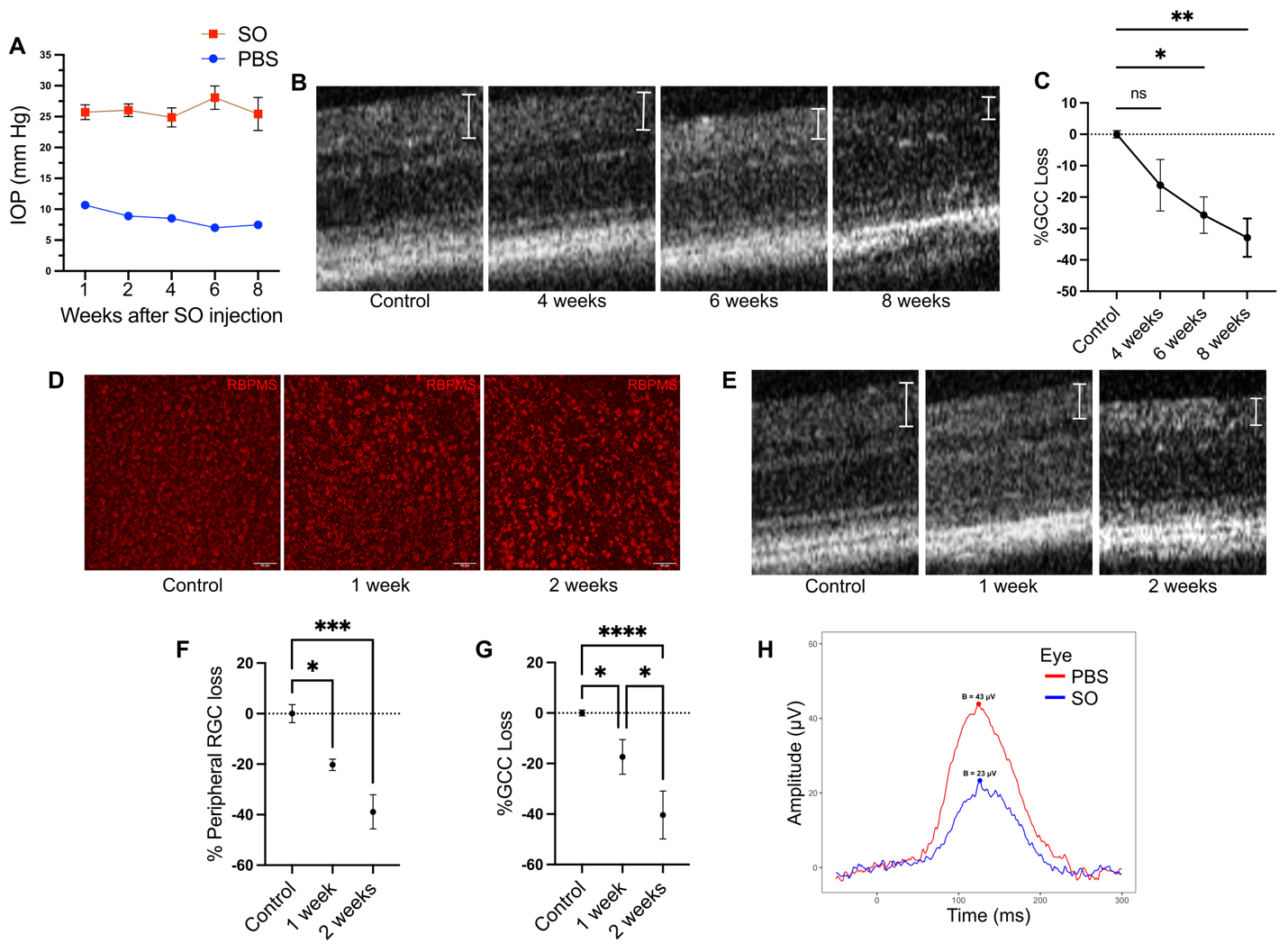

**Supplementary Figure 1: Silicon oil-based moderate and severe models of OHT. (A)** IOP measurement at different time points (1, 2, 4, 6, 8 wks) for moderate OHT (n=20). (**B)** OCT images of the retinal layers depicting retinal thinning over the time course of moderate OHT. Ganglion cell complex (GCC) thickness is marked by line segments (**C)** Quantification of ganglion cell layer measurements represented as percent GCC loss compared to mean in normotensive retinas. Data were analyzed by One-way ANOVA with Tukey’s multiple comparisons test (*p<0.05; **p<0.01). (**D)** Representative confocal micrographs of RGCs stained for RNA binding protein (RBPMS) at different time points of severe OHT. (**E)** OCT images of retinal layers depicting retinal thinning over the course of severe OHT. Ganglion cell complex (GCC) thickness is marked by bar arrows. (**F)** Percent loss of peripheral RGCs at different time points of severe OHT compared to normotensive controls. Data were analyzed by One-way ANOVA with Tukey’s multiple comparison test (*p<0.05; ***p<0.001). (**G)** Quantification of the GCC layer thickness. Data was analyzed by One-way ANOVA with Tukey’s multiple comparisons test (*p<0.05; ****p<0.0001; ns, not significant). (**H)** Representative p-STR recording showing a reduction in b-wave amplitude after 2 wks of severe OHT. Data presented as mean±SEM (n=4-6 for B-H**).**

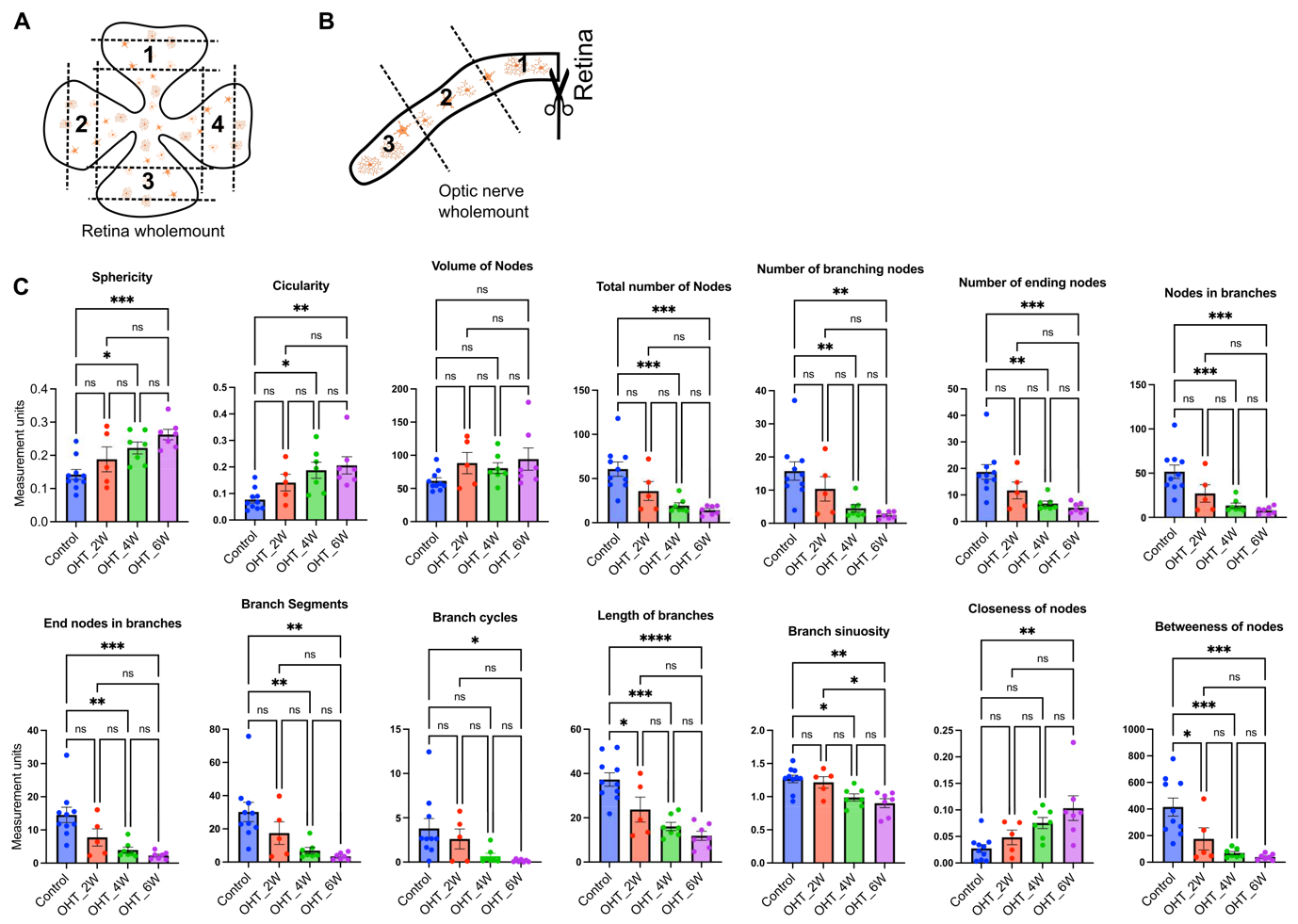

**Supplementary Figure 2: Dynamic morphological changes in retinal microglia during moderate OHT. (A)** Schematics showing confocal imaging locations in the retinal wholemount. **(B)** Schematics showing section of optic nerve that was selected for optic nerve wholemount and confocal imaging. **(C)** Specific features (number of features=14) of microglia morphology analyzed by a feature extraction tool for normotensive control (n=10) and 2 wks (n=5), 4 wks (n=7), and 6 wks (n=7) of moderate OHT. Each dot represents the mean value of a feature from 1 retina. Data was analyzed by One-way ANOVA with Tukey’s multiple comparison test (*p<0.05; **p<0.01; ***p<0.001; ****p<0.0001; ns, not significant). Data presented as mean±SEM.

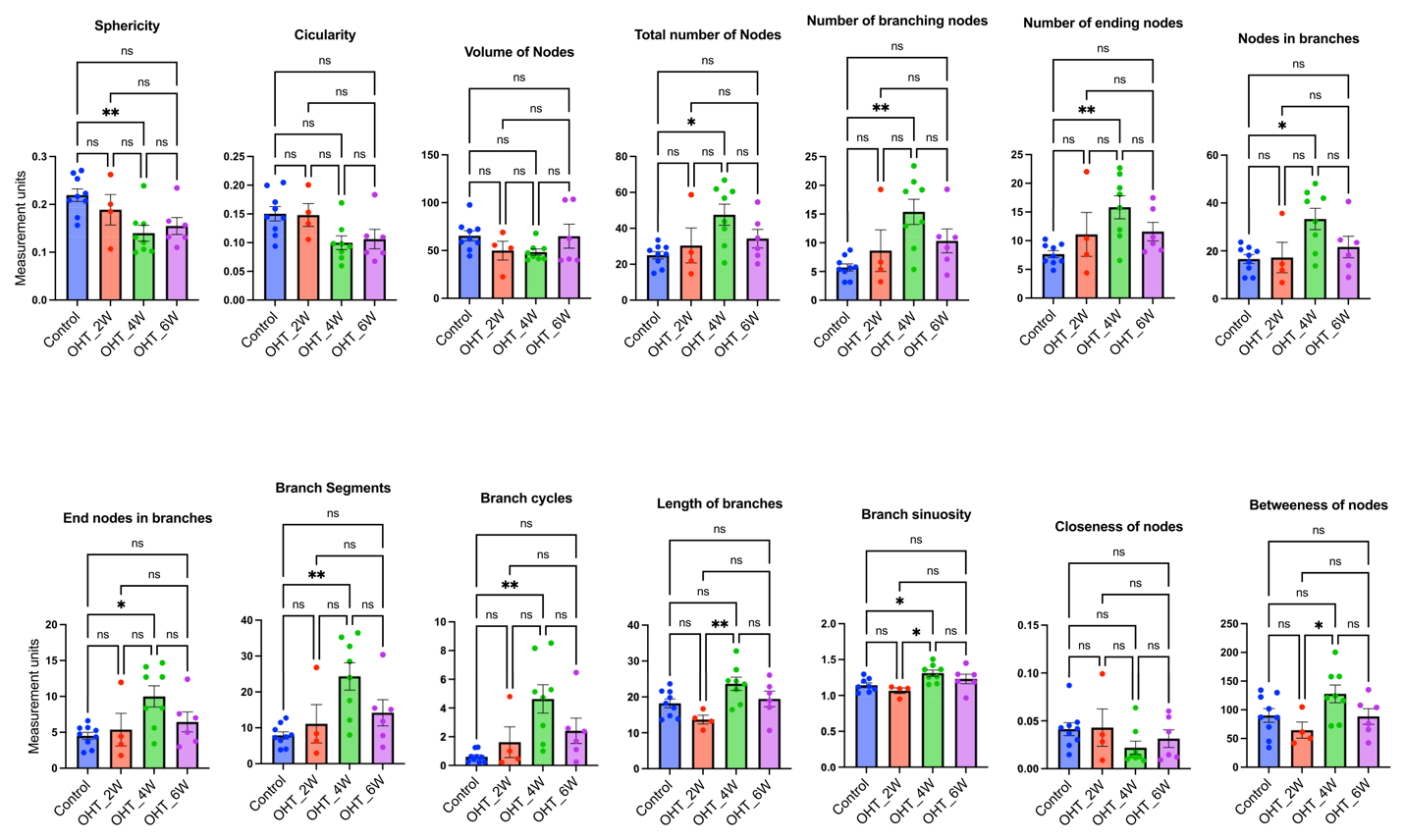

**Supplementary Figure 3: Morphological response of optic nerve microglia to moderate OHT.** Specific features (number of features=14) of microglia morphology analyzed by the feature extraction tool for normotensive control (n=9) and 2 wks (n=4), 4 wks (n=8), and 6 wks (n=6) of moderate OHT. Each dot represents the mean value of the feature from 1 retina. Data was analyzed by One-way ANOVA with Tukey’s multiple comparison test (*p<0.05; **p<0.01; ns, not significant). Data presented as mean±SEM.

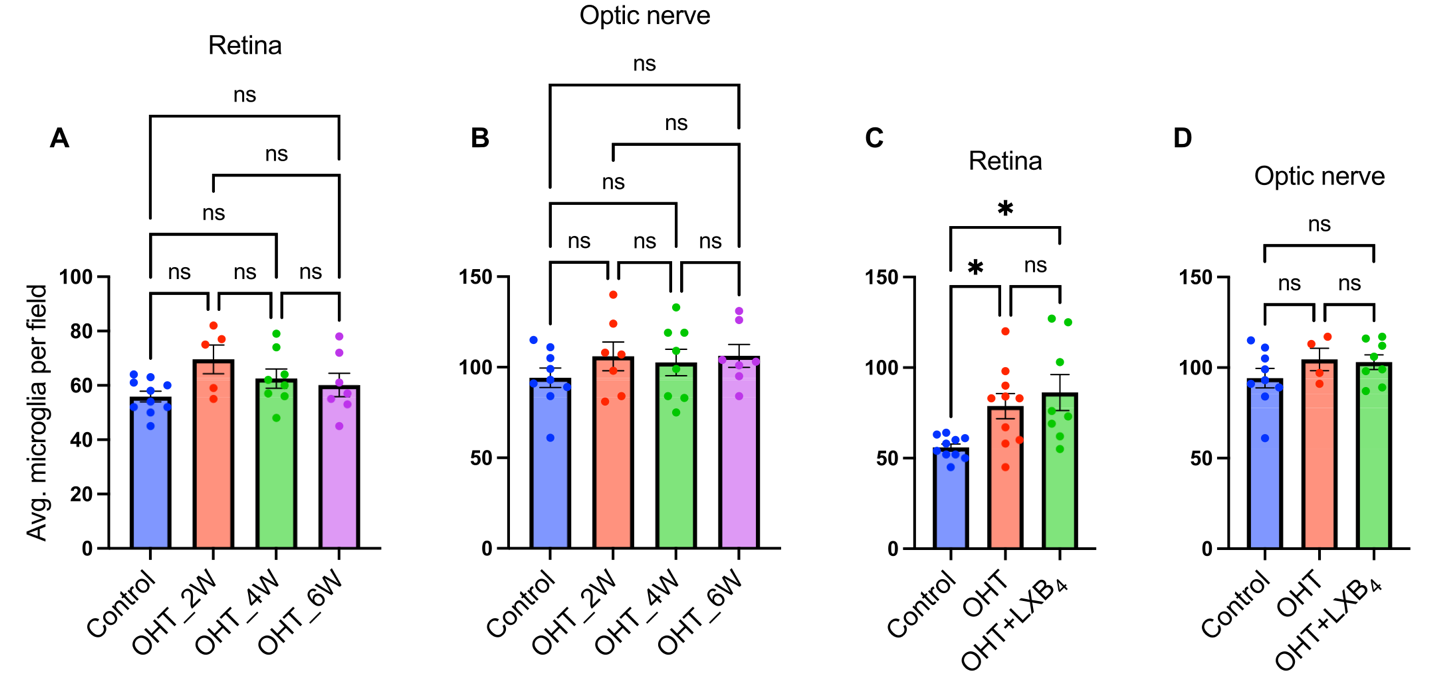

**Supplementary Figure 4: Microglia counts in the retina and optic nerve in response to moderate and severe OHT. (A)** Microglia counts in the retinal wholemounts in response to moderate OHT at 2 wks (n=5), 4 wks (n=8), and 6 wks (n=7). **(B)** Microglia counts in the optic nerve wholemounts in response to moderate OHT at 2 wks (n=7), 4 wks (n=8), and 6 wks (n=7). **(C)** Microglia counts in the retinal wholemounts in response to severe OHT with or without LXB_4_ treatment at 1wk. Normotensive control (n=10), OHT (n=10), and OHT+LXB_4_ (n=8). **(D)** Microglia counts in the optic nerve wholemounts in response to severe OHT with or without LXB_4_ treatment at 1wk. Normotensive control (n=9), OHT (n=4), and OHT+LXB_4_ (n=8). Each dot represents the mean count from 1 retina and optic nerve. Data was analyzed by One-way ANOVA with Tukey’s multiple comparison test (*p<0.05; ns, not significant). Data presented as mean±SEM.

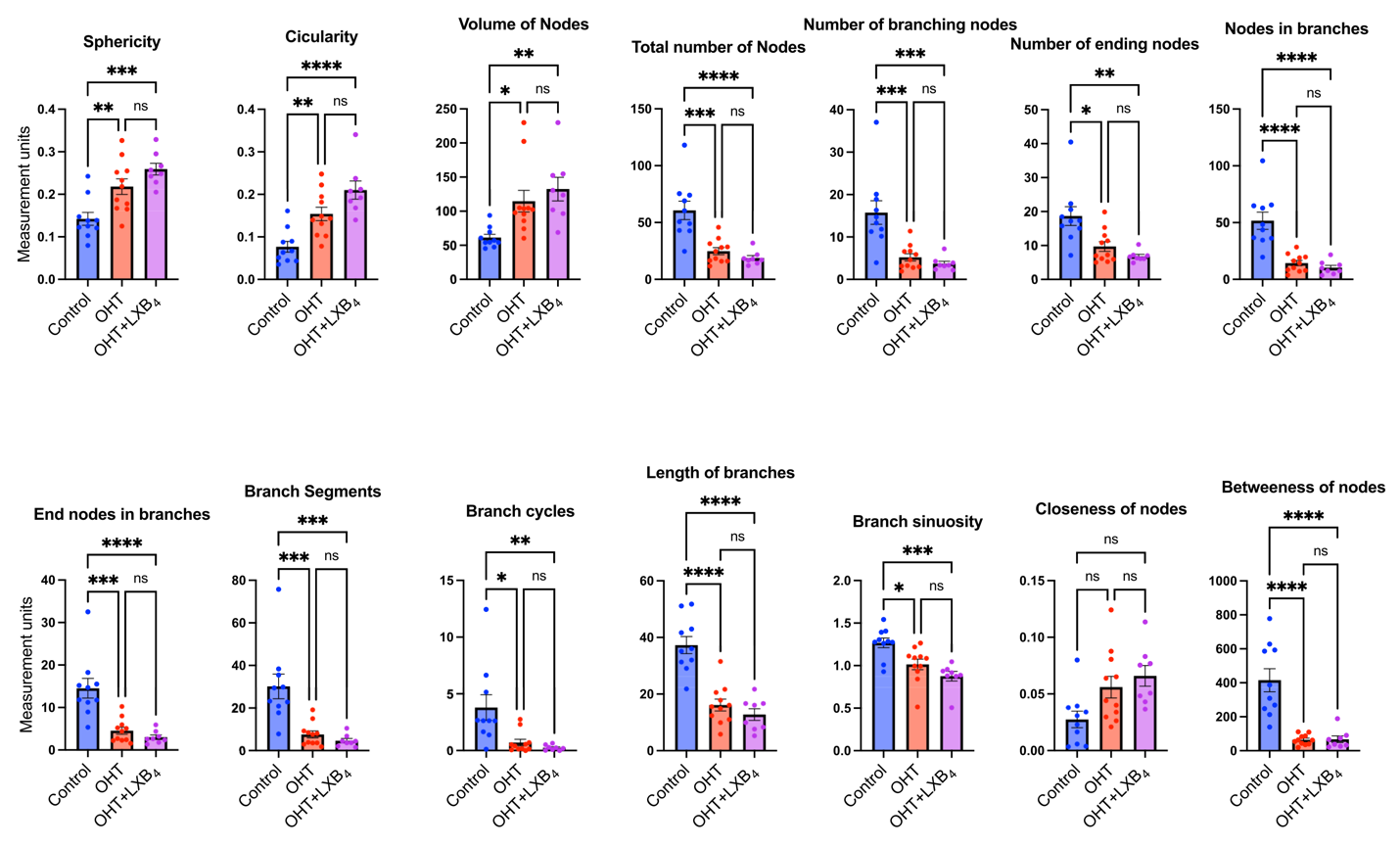

**Supplementary Figure 5: Dynamic morphological changes in retinal microglia during severe OHT and LXB_4_ treatment.** Specific features (number of features=14) of microglia morphology analyzed by a feature extraction tool for normotensive control (n=10), OHT (n=11), and OHT+LXB_4_ (n=8). Each dot represents the mean value of a feature from 1 retina. Data was analyzed by One-way ANOVA with Tukey’s multiple comparison test (*p<0.05; **p<0.01; ***p<0.001; ****p<0.0001; ns, not significant). Data presented as mean±SEM.

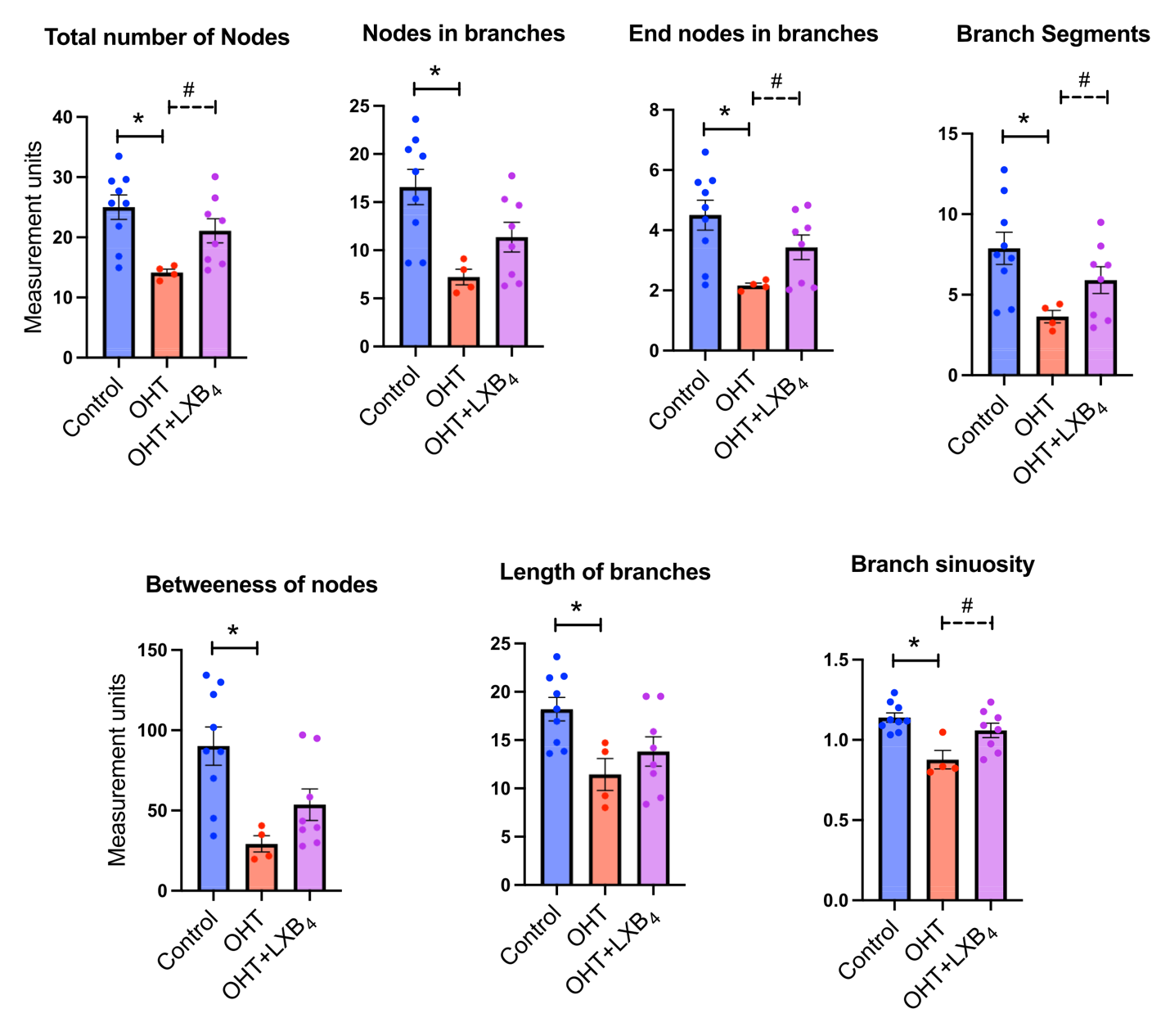

**Supplementary Figure 6: Morphological changes in optic nerve microglia during severe OHT and LXB_4_ treatment.** Specific features (number of features=7) of microglia morphology analyzed by a feature extraction tool for normotensive control (n=9), OHT (n=4), and OHT+LXB_4_ (n=8). Each dot represents the mean value of a feature from 1 optic nerve. Data was analyzed by One-way ANOVA with Tukey’s multiple comparison test (*p<0.05) and unpaired t-test (#p<0.05) between OHT+LXB_4_ versus OHT groups. Data presented as mean±SEM.

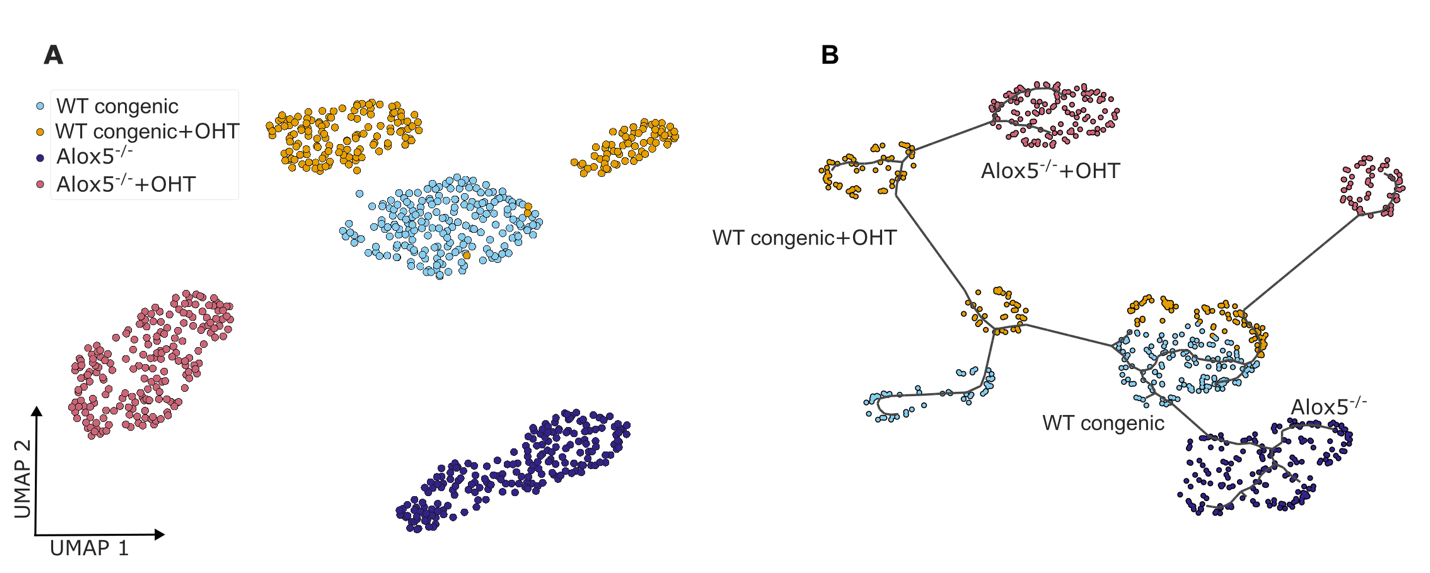

**Supplementary Figure 7: Lipoxin deficiency correlates with an altered optic nerve microglia phenotype. (A)** morphOMICs analysis of *Alox5*^-/-^ and congenic wild-type control (*Alox5^+/+^)* optic nerve microglia in normotensive eyes and after 1 wk of severe OHT. **(B)** Pseudotime trajectory analysis of optic nerve microglia populations from *Alox5*^-/-^ normotensive eyes and after 1 wk of severe OHT, optic nerve microglia from normotensive congenic *Alox5^+/+^* are displayed as the root cells.

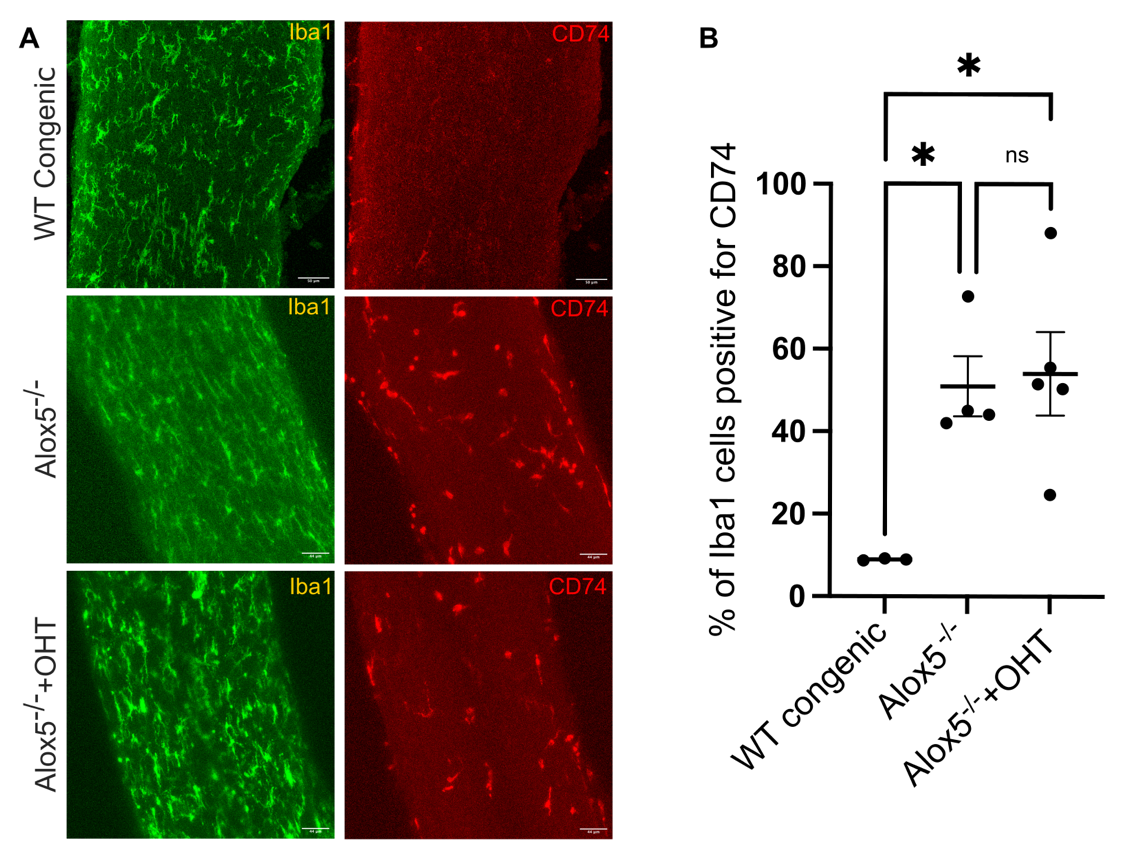

**Supplementary Figure 8: Lipoxin deficiency correlates with the presence of a CD74 positive disease -associated microglia population in the optic nerve. (A)** Micrographs of Iba1 (green) and CD74 (red) stained optic nerve sections from WT normotensive congenic, Alox5^-/-^ normotensive and after 1 wk of severe OHT (scale bar - 44 μm). **(B)** Quantification of CD74 and Iba1 positive cells. Data was analyzed by One-way ANOVA with Tukey’s multiple comparison test (*p<0.05; ns, not significant). Data presented as mean±SEM.

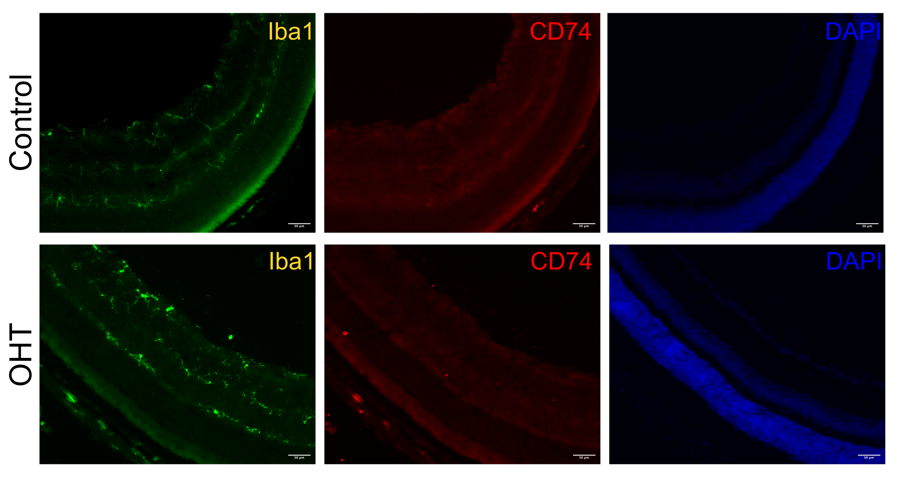

**Supplementary Figure 9: CD74 positive disease-associated microglia are not present or induced in the retina in normotensive or hypertensive eyes.** Micrographs of retinal sections stained with Iba1 (488 nm) and CD74 (594 nm) along with DAPI (405 nm) in normotensive control and severe OHT (scale bar- 50μm).

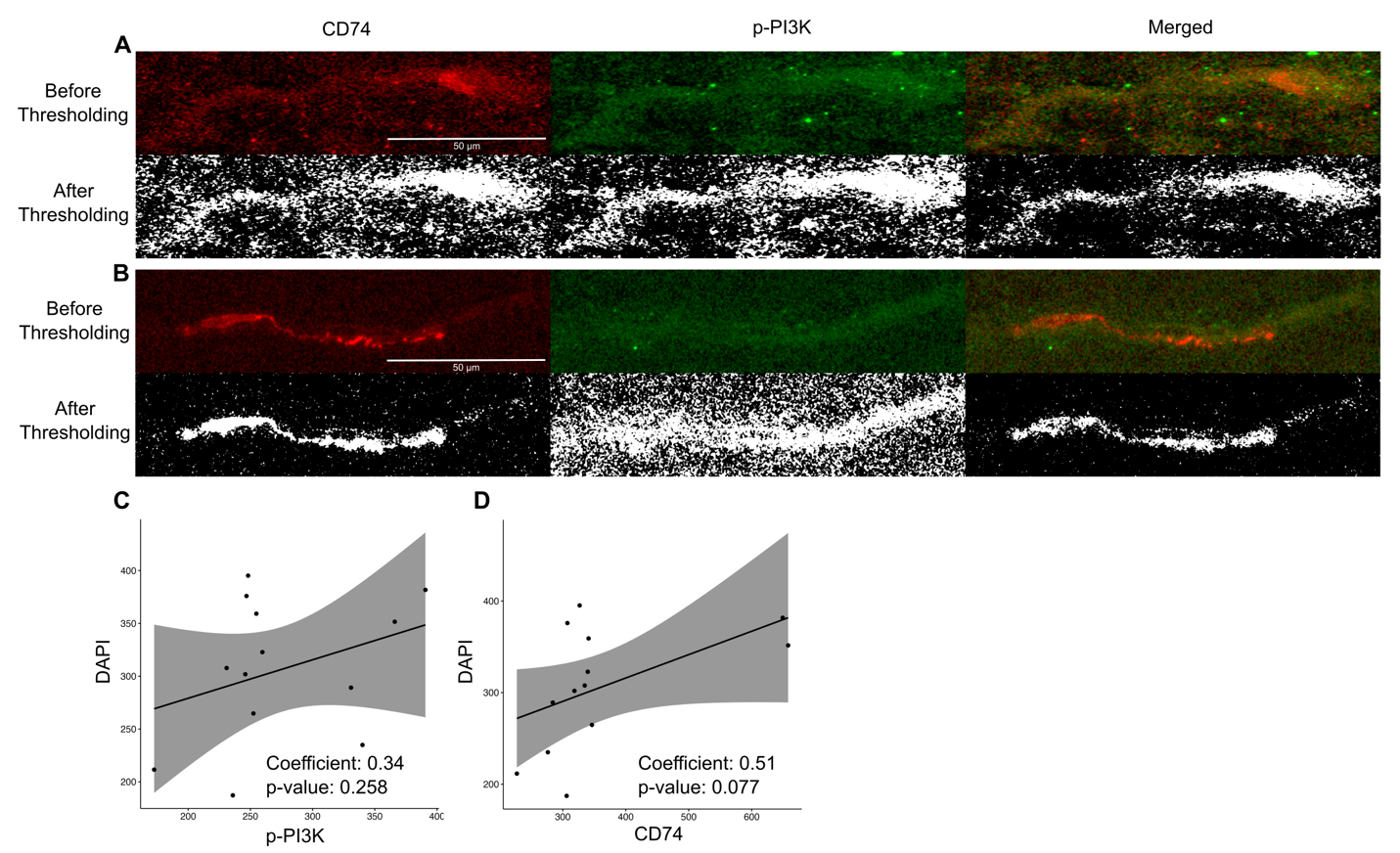

**Supplementary Figure 10: Colocalization of CD74 and p-PI3K in the optic nerve sections. (A, B) (A)** Representative BIOP analysis output for CD74 and p-PI3K colocalization in OHT and **(B)** OHT+LXB_4_**.** (**D)** Correlation analysis for DAPI and p-PI3K and (**F)** DAPI and CD74 colocalization after 1 wk of severe OHT.
